# Cell-adaptable dynamic hydrogel reinforced with stem cells improves the functional repair of spinal cord injury by alleviating neuroinflammation

**DOI:** 10.1101/2021.09.24.461661

**Authors:** Xin Yuan, Weihao Yuan, Lu Ding, Ming Shi, Liang Luo, Yong Wan, Jiwon Oh, Yanfang Zhou, Liming Bian, David Y.B. Deng

## Abstract

Spinal cord injury (SCI) is one of the most challenging clinical issues. It is characterized by the disruption of neural circuitry and connectivity, resulting in neurological disability. Adipose-derived stem cells (ADSCs) serve as a promising source of therapeutic cells for SCI treatment. However, the therapeutic outcomes of direct ADSCs transplantation are limited in the presence of an inflammatory microenvironment. Herein, a cell-adaptable neurogenic (CaNeu) hydrogel was developed as a delivery vehicle for ADSCs to promote neuronal regeneration after SCI. The dynamic network of CaNeu hydrogel loaded with ADSCs provides a cell-infiltratable matrix that enhances axonal growth and eventually leads to improved motor evoked potential, hindlimb strength, and coordination of complete spinal cord transection in rats. Furthermore, the CaNeu hydrogel also establishes an anti-inflammatory microenvironment by inducing a shift in the polarization of the recruited macrophages toward the pro-regeneration (M2) phenotype. Our study showed that the CaNeu-hydrogel‒mediated ADSCs delivery resulted in significantly suppressed neuroinflammation and apoptosis, and that this phenomenon involved the PI3K/Akt signaling pathway. Our findings indicate that the CaNeu hydrogel is a valuable delivery vehicle to assist stem cell therapy for SCI, providing a promising strategy for central nervous system diseases.

## 1. Introduction

Spinal cord injury (SCI) is one of the most challenging clinical issues with complex pathological condition, leading to variable degrees of permanent neurological dysfunction [1, 2]. Current therapeutic interventions targeting SCI include hemodynamic monitoring in the intensive care unit, early surgical decompression, blood pressure augmentation, and administration of methylprednisolone; however, the efficacy of these treatment strategies is limited [3]. Hence, transplantation of stem cells has been investigated as a potential therapy for SCI to promote the regeneration of injured axons [4–6]. The use of adipose-derived stem cells (ADSCs) has several advantages, such as abundant availability, low immunogenicity, excellent expansion potential, and requirement of less invasive techniques for cell collection [7, 8]. Several preclinical studies have demonstrated the potential ability of ADSCs to repair SCI in a paracrine manner [9–11]. In addition, a clinical trial investigating the use of autologous ADSCs for treating patients with cervical SCI is currently underway at the Mayo Clinic [12]. The results of this present study suggest that ADSCs administration can promote neural tissue preservation immediately after SCI and eventually promote functional recovery.

In clinical trials, direct local transplantation of stem cells into the central lesion of SCI typically results in extensive cell loss and death owing to the development of severe inflammatory microenvironment after injury [13]. Cystic cavities, complicated by the development of traumatic or vascular injuries, can further aggravate cell loss [13]. Furthermore, inflammation post-SCI contributes to the formation of a glial scar, which acts as a chronic and physical barrier that prevents axonal regeneration [14, 15]. Therefore, biomaterial scaffolds hold considerable promise with respect to enhancing the efficacy of SCI repair [16–19], and hydrogels rank among the most ideal carriers for stem cell delivery. Previous studies have shown that hydrogels, including hyaluronic acid (HA) [20–22], poly(lactic-co-glycolic acid) (PLGA) [23, 24], poly (2-hydroxyethylmethacrylate) (HEMA) [25, 26], nanofiber [27–29], and self-assembled peptide hydrogels [30–32], can be engineered to mimic the architecture of the lost ECM in the spinal cord and can structurally support cell migration and axonal regrowth. Our previous study reported a Pluronic F-127 (PF-127)-based thermo-sensitive hydrogel for enhancing SCI treatment [33]. However, most of the existing hydrogels cannot completely recapitulate the dynamic properties of natural ECM, which are critical for supporting various cellular activities essential for neural regeneration, including migration, proliferation, differentiation, and axonal growth. However, hydrogels with a cell-adaptable dynamic network would better support the expansion and differentiation of encapsulated stem cells and recruitment of host immune cells than hydrogels with encapsulated stem cells alone. Therefore, we sought to develop a cell-adaptable hydrogel for ADSCs to promote the neurogenesis and subsequent neuronal relay formation after SCI.

In this study, we aimed to develop a structurally dynamic supramolecular CaNeu hydrogel based on “Host–guest” crosslinking to mimic the structural and mechanical properties of neural tissues. We anticipate that the dynamic network of the CaNeu hydrogel will provide a favorable environment for neural differentiation and axonal growth of encapsulated ADSCs to promote nerve regeneration. Further, we investigated the effect of the ADSCs-loaded CaNeu hydrogel on M2 macrophages to evaluate SCI healing in an animal model during eight weeks **(Scheme 1)**.

**Scheme 1.**
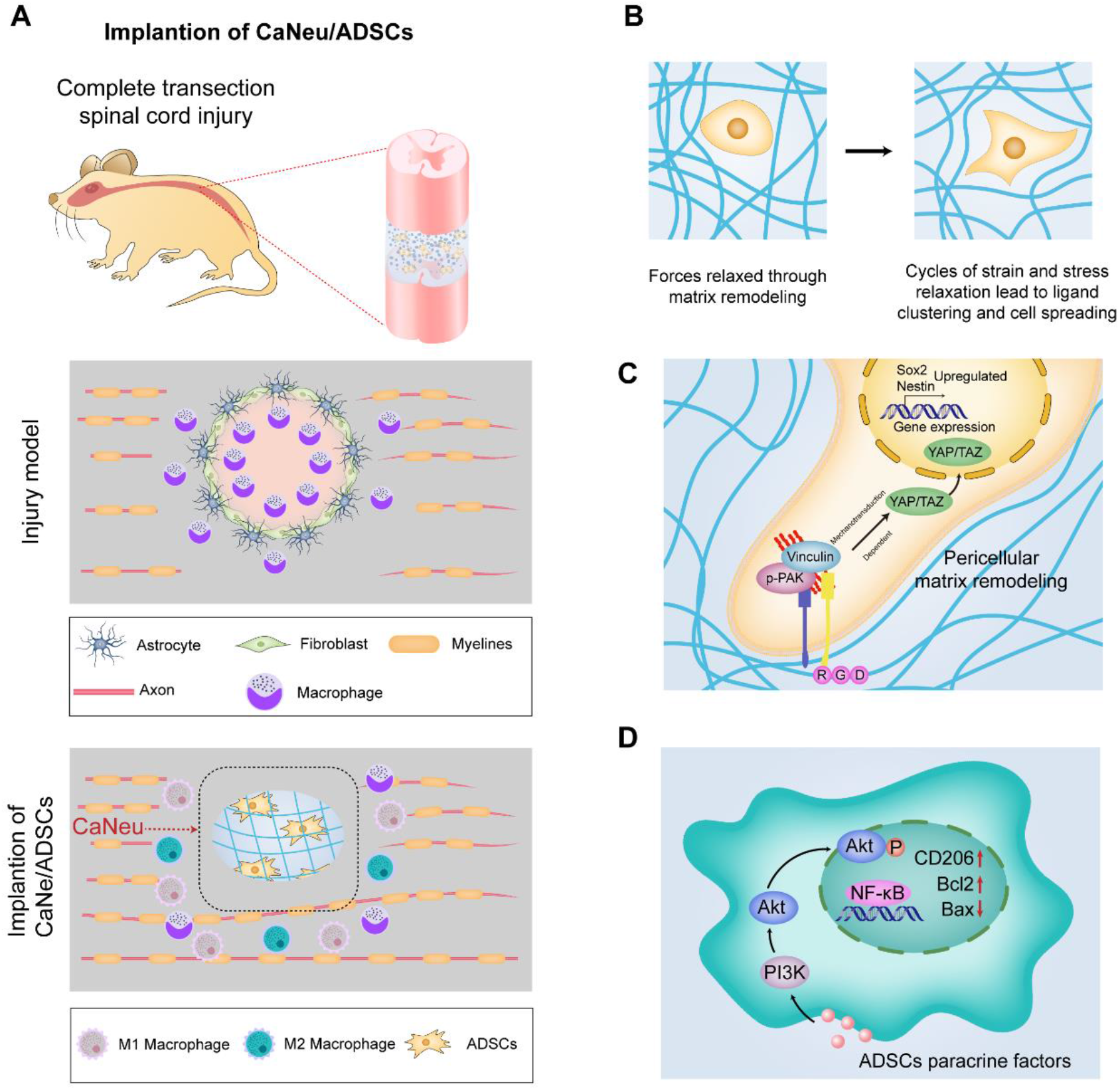
Schematic illustration depicting the stemness maintenance and promotion of M2 macrophage polarization by ADSCs-loaded dynamic CaNeu hydrogel. (A) A proposed strategy for the effective modulation of SCI microenvironment after injuries by developing a ADSCs-loaded dynamic CaNeu hydrogel-based therapeutic interventions. (B) Matrix remodeling of dynamic CaNeu hydrogel promotes spreading of ADSCs. (C) Schematic depicting a mechanism by which increased cell spreading facilitated by soft dynamic CaNeu hydrogel regulates YAP expression and stemness property. (D) CaNeu hydrogel dynamic matrix encapsulated ADSCs enhanced functional recovery of SCI rats by promotion of M2 macrophage polarization. ADSC induces activation of PI3K/Akt pathway through paracrine factors, leading to the phosphorylation of the Akt and the change of Akt expression level results in an increase of M2 macrophages. Furthermore, apoptosis was suppressed by upregulation of Bcl-2 and downregulation of Bax. These events promote the spreading of ADSCs and alleviate neuroinflammation following SCI in the niche microenvironment, thereby promoting SCI repair.

## 2. Materials and methods

### 2.1 Synthesis of CaNeu and GelMA

β-CD (10 g) was dissolved in 150 mL dimethylformamide (DMF), followed by the addition of 3.5 mL triethanolamine (TEA) to the solution. The mixture was cooled to 0℃ and 2.5 mL acryloyl chloride was added dropwise into the solution for 1 h. The mixture was filtered to remove the precipitated TEA-HCl salt, and the obtained clear yellowish solution was concentrated to 20 mL using rotary evaporation under vacuum. The solution was poured into 800 mL acetone to obtain an Ac-β-CD precipitate, and the precipitate was washed with acetone three times. Ten grams of gelatin (type A) was dissolved in 100 mL of PBS at 50℃. Methacrylic anhydride (12 mL) was then added to the 10% gelatin solution, and the reaction was incubated at 50℃ for 4 h. The CaNeu hydrogel network stabilized by the reversible “host-guest” complexes was formed by the UV-mediated polymerization of acryloyl group in Ac-β-CD. The resulting mixture was dialyzed against deionized (DI) water for one week at 45℃. The dry product was obtained by lyophilization for three days. The substitution degree of Ac-β-CD in CaNeu and the modification rate of GelMA were determined using ^1^H nuclear magnetic resonance (^1^H NMR) at 400 MHz (Bruker, USA) in DMSO-*d*_6_ and D_2_O at 25℃, respectively **(Figure S1)**.

### 2.2 Rheological and uniaxial compression tests

All rheological and uniaxial compression tests were performed on Kinexus Lab Plus (Malvern, UK). For the oscillatory time sweep, we set the following parameters: frequency sweep (0.01–10 Hz), strain sweep (0.1%–1000%), alternating high and low strain sweep (1% and 600%, respectively), and gap between rotor and plate to 0.5 mm. For the uniaxial compression tests, the prefabricated hydrogels (ϕ = 5 mm, h = 2 mm) were located at the center of the plate, and the compression rate was set at 0.05 mm/s. Young’s modulus was calculated from the initial linear region of the stress–strain curves (strain < 10%). The stress relaxation test was conducted using a general sequence from the instrument manufacturer with an initial compression of 20%.

### 2.3 Scanning electron microscopy (SEM)

The hydrogels were frozen by immersion in liquid nitrogen for 5 min, followed by lyophilization for three days. The completely dried hydrogels were cut into small pieces to investigate the internal morphology. SEM was conducted using the SU8010 ultra-high resolution SEM (Hitachi High Technology, Japan) with an accelerating voltage of 5 kV.

### 2.4 Model protein (BSA) release

BSA was mixed with the precursor solutions of CaNeu or GelMA hydrogels before gelation as described above (100 μg BSA for each hydrogel). The prepared hydrogels (n = 6 for each group) were immersed in 1 mL PBS containing 0.1% (w/v) NaN_3_. Fifty microliters of PBS from each sample was collected from the supernatant on days 0, 1, 3, 7, 14, and 28. The BSA content was quantified using a BCA protein quantification kit and the percentage of released BSA was calculated using the following equation:

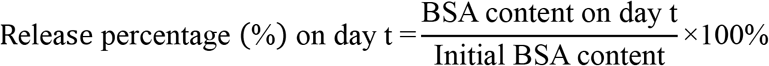

### 2.5 Hydrogel degradation test

Degradation test: The prepared hydrogels (n = 6 for each group) were immersed in 1 mL PBS containing 0.1% (w/v) NaN_3_. The samples were removed and washed thrice with DI water to eliminate residual salts. Further, the samples were immersed in liquid nitrogen for 5 min and lyophilized for three days. The dry mass of each sample at different time points was measured, and the weight percentage was calculated using the following equation:

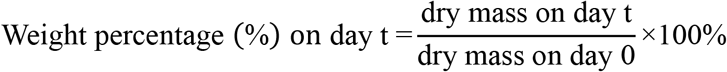

### 2.6 Preparation and identification of ADSCs

ADSCs were isolated from rat epididymal fat tissue using a standard procedure [34]. After washing with 10% and 5% penicillin–streptomycin solution (Gibco), the tissue was cut into pieces and enzymatically digested using 1 mL 0.5% type I collagenase (Sigma) at 37°C for 60 min under shaking conditions. The mixture was centrifuged at 300 × *g* for 5 min and resuspended in the medium three times. Obtained cells were resuspended in Dulbecco’s modified Eagle’s medium (DMEM; Gibco) containing 10% fetal bovine serum (FBS; Gibco) and cultured at 37°C in 5% CO_2_. ADSCs at passages 3–5 were prepared for subsequent experiments.

### 2.7 Viability and proliferation of encapsulated ADSCs in CaNeu hydrogel

For *in vitro* cell experiments, ADSCs were suspended in GelMA and CaNeu hydrogels at a density of 5 × 10^6^ cells mL^−1^, crosslinked, and cultured in DMEM/FBS medium at 37°C. For longer culture conditions, 1% PEGDA was added, so as not to affect the properties of the CaNeu hydrogel **(Figure S2)** [35].

Cell infiltration test was done using the 24-well transwell. GelMA and CaNeu hydrogels with volume of 20 μL were formed on the top of the transwell membrane, and then the transwell was placed into a 24-well plate. 50 μL of ADSC suspension (5 × 10^6^ cells mL^−1^) in basal medium was added to the top of the hydrogel, and 980 μL growth media with 10 ng mL^−1^ SDF-1α was added into each well. After 12 h of incubation, the hydrogels were fixed by 4% paraformaldehyde for 30 min and stained with DAPI. The acquired confocal microscope images were reconstructed for three-dimensional visualization of the infiltrated cells in the hydrogels. Cell viability on day 1 and day 14 was evaluated using the LIVE/DEAD™ cell imaging kit (Thermo Fisher Scientific) as per the manufacturer’s instructions. Hydrogel samples were added to 100 μL of live/dead mixture and incubated at 25°C for 20 min. Live/dead and infiltration assays were performed, and cells were observed under a confocal microscope (Zeiss LSM 880, Germany). Morphology of ADSCs was quantified using the F-factor (F = 4πA/P^2, where A = area and P = perimeter).

### 2.8 Biochemical analysis of stemness and differentiation markers

To investigate the biochemistry of ADSCs in GelMA or CaNeu, pcDNA3.1-Sox2 plasmids were constructed and transfected using Lipofectamine 3000 (Thermo Fisher Scientific).The expression levels of *nestin* (forward primer: 5′-CTCTGGGCAAGTGGAACG-3′; reverse primer: 5′-TCCCACCGCTGTTGATTT-3′), *Sox2* (forward primer: 5′-GCGGAGTGGAAACTTTTGTCC-3′; reverse primer: 5′-CGGGAAGCGTGTACTTATCCTT-3′), Vimentin (forward primer: 5′-CCGCTTCGCCAACTACAT-3′; reverse primer: 5′-CGCAACTCCCTCATCTCCT-3′), and YAP (forward primer: 5′-GATGGATGGGAGCAAG-3′; reverse primer: 5′-GCAAAACGAGGGTCAA-3′) were measured using quantitative reverse transcription polymerase chain reaction (qRT-PCR). ADSCs located in GelMA or CaNeu were lysed using Trizol (Thermo Fisher Scientific), and RNA was isolated using phenol– chloroform extraction. RNA (1 μg) was reverse transcribed using the Perfect real-time cDNA reverse transcription kit (TaKaRa). Quantitative PCR was performed with 1 μg of cDNA per sample per target gene using AceQ qPCR SYBR Green master mix (Vazyme) on a CFX 96 Touch real-time PCR system (Bio-Rad). For YAP-TEAD inhibition studies, ADSCs in GelMA or CaNeu were cultured in the presence of 5 × 10^−7^ M verteporfin.

To induce neuronal differentiation, the culture medium was replaced with Neurobasal (Invitrogen), 2% B27 (Gibco), and 10 ng mL^−1^ FGF-2 (PeproTech). For immunocytochemistry, samples were fixed with 4% paraformaldehyde, permeabilized with 0.2% Triton X-100, and blocked with 5% BSA. Samples were probed overnight with primary antibody against chicken MAP2 (1:5000; Abcam), rabbit NeuN (1:200; Abcam) at 4°C. The samples were washed and subsequently with secondary antibodies against Alexa Fluor 647 (1:400, Abcam), followed by DAPI staining for 2 h at 25℃. The samples were washed and imaged using a Zeiss LSM880 confocal microscope, and the results were reconstructed in three dimensions using Image J (NIH) [36].

### 2.9 Establishment of complete transection of spinal cord animal model

All animal experimental procedures were performed in compliance with the guidelines proposed by the China Council on Animal Care and were approved by the Institutional Animal Care and User Committee of Guangdong Medical University, Dongguan, China.

For animal experiments, approximately 10 μL of the hydrogels were crosslinked in a polytetrafluoroethylene (PTFE) mold (diameter, 2 mm; depth, 2 mm). Further, ADSCs (1 × 10^7^ cells mL^-1^) were loaded onto CaNeu hydrogels. Adult female Sprague Dawley rats (200–220 g, n = 89) from Southern Medical University, Guangzhou of China were used in this study. The animals were deeply anesthetized with intraperitoneal injections of sodium pentobarbital (3%, 50 mg kg^−1^). Dorsal laminectomy was performed at the 10^th^ thoracic (T9–T10) vertebrae to expose the spinal cord [37, 38]. Further, the T9–T10 spinal cord tissue (2 mm) was excised, resulting in a total cross-sectional injury. Scaffold transplantation or cell injection (1 × 10^7^ cells mL^−1^, 10 μL) was followed by layer-by-layer sutures. All animals were randomly divided into five groups containing CaNeu/ADSCs (n = 19), CaNeu (n = 19), ADSCs (1 × 10^7^ cells mL^−1^, n = 19), SCI (no transplantation, n = 19), and sham (animals received laminectomy and dura opening but no spinal cord transection, n = 13). After surgery, all animals were injected with penicillin for seven days and artificial urination was employed every day until partially autonomous urination was restored.

### 2.10 Perfusion and tissue processing

The animals were deeply anesthetized with sodium pentobarbital (3%, 50 mg kg^−1^) and perfused through the left ventricle with saline and 4% paraformaldehyde for fixation. The dissected spinal cords **(Figure S7A)** were fixed in PFA for 6 h and incubated in PBS containing 30% sucrose at 4°C until the tissues sank. All samples were embedded in Tissue-Tek O.C.T (Sakura, USA) at −30°C and sectioned using a cryostat (Thermo Fisher Scientific). H&E staining of spinal cord sections was performed in accordance with standard staining protocols. The animals were euthanized 1 week, 2weeks, and 8 weeks post injury, depending on the experiment performed **(Table S1)**.

### 2.11 Immunofluorescence

The spinal cord slices were blocked with 5% BSA prepared in PBS containing Tween 20 for 2 h at room temperature, and probed overnight at 4°C with the following primary antibodies: rabbit anti-TUBB3 (1:200; CST), mouse anti-GFAP (1:200; CST), rabbit anti-5HT (1:10,000; Immunostar), rabbit anti-NF200 (1:50; CST), mouse anti-S100β (1:100; CST), rabbit anti-MBP (1:50; CST), cleaved caspase-3 (1:500; CST), rabbit anti-IBA1 (1:50; CST), mouse anti-CD68 (1:100; Abcam), and rabbit anti-CD163 (1:100; Thermo Fisher Scientific). The sections were then washed and incubated with Alexa Fluor 488- or 594-conjugated secondary antibodies (1:500; Thermo Fisher Scientific) for 2 h. Nuclei were stained with DAPI (1:2000; Sigma-Aldrich) for 10 min. The stained sections were examined using a Zeiss LSM 880 confocal microscope.

### 2.12 Western blotting

For western blotting, we used standard western blotting techniques and BeyoECL Plus chemiluminescence kit (Beyotime). The following antibodies were used: cleaved caspase-3 (1:1000; CST), IL-6 (1:1,000; Affinity), Akt (1:1,000; CST), pAkt (1:1,1000; CST), Bcl-2 (1:1,000; Affinity), Bax (1:1,000; Affinity), and GAPDH (1:1,000; Beyotime). Densitometry was performed using ImageJ (NIH), and the expression of target proteins was normalized to GAPDH.

### 2.13 Functional analysis

#### 2.13.1 Electrophysiology analysis

MEP of the different groups was examined on week 8. After the rats were anesthetized, a stimulation electrode was inserted into the rostral extremity of the surgically exposed spinal cord region, and the recording electrode was inserted into the biceps flexor cruris to record the response after simulation. The amplitude was measured from the initiation point of the first response wave to its highest point. The peak-to-peak amplitude (P-P value) was calculated to estimate the nerve conduction function in the hindlimb.

#### 2.13.2 Behavioral assessment

Behavior analyses were performed for rats subjected to spinal cord complete transection based on the results of BBB open field locomotor test [39]. The BBB score for the evaluation of hindlimb function after SCI ranged from 0 (flaccid paralysis) to 21 (normal gait). The animals were allowed to walk around freely in an open field for 3 min, and hindlimb movements were closely observed. Recovery of the hindlimb for each group was assessed once a week for eight weeks. The locomotor functions of the hindlimb were scored in accordance with the BBB scoring system, including the frequency and quality of hindlimb movement as well as forelimb/hindlimb coordination.

#### 2.13.3 Morphometric evaluation and ultrastructure analysis

Eight weeks after treatment, the animals (n = 3 per group) were anesthetized with pentobarbital sodium as described above, and perfused transcardially with 4% paraformaldehyde and 2% glutaraldehyde in 0.1 M phosphate buffer (pH 7.4). The lesion epicenter was extracted and post-fixed by immersion in 1% osmium tetroxide at 4°C for 10 min, followed by incubation with 1.5% potassium ferrocyanide in 0.1 M cacodylate buffer for 1 h at 4°C. Further, the sections were stained with 2% uranyl acetate in 30% ethanol for 30 min at 4°C. Lastly, the sections were dehydrated in a graded acetone series of ethanol at 4°C. The sections were rinsed with propylene oxide (3 × 20 min) at 25°C prior to embedding, followed by 100% Spurr’s resin for 5–6 h. The resin was polymerized for a minimum of 16 h at 60°C in plastic molds until it hardened into resin blocks. Semi-thin (500 nm) and ultrathin (70 nm) sections from each group were cut on an RMC ultramicrotome (USA). The semithin sections were stained with toluidine blue and imaged under a Leica DM4B microscope (Germany).

### 2.14 Statistical analysis

All data are shown as the mean ± standard error of the mean (SEM). All statistical analyses were performed using GraphPad Prism 8.2, with two-tailed unpaired Student’s *t*-test for comparison between two groups or one-way analysis of variance (ANOVA) with Bonferroni’s or Tukey’s post-hoc test for comparing multiple groups. We performed Tukey’s multiple comparison test in two-way ANOVA for the BBB score. *, **, ***and **** have been used to indicate *P* < 0.05, *P* < 0.01, *P* < 0.01, and *P* < 0.0001, respectively.

## 3. Results

### 3.1 The CaNeu hydrogel exhibits various dynamic properties

We examined the dynamic properties of CaNeu hydrogel stabilized by reversible host–guest crosslinks and compared them with those of the conventional methacrylated gelatin (GelMA) hydrogel crosslinked by irreversible covalent bonds **(Figure 1A)**. To mimic the stiffness of the spinal cord (500–1500 Pa [40]), the stiffness of the CaNeu hydrogel was controlled at ~900 Pa. The stiffness of the GelMA hydrogel was also controlled at ~900 Pa to decouple the influence of hydrogel stiffness from that of hydrogel dynamics **(Figure S3A)**. The storage modulus (G’) of the CaNeu hydrogel exhibited significant frequency-dependent behavior, whereas the G’ of the GelMA hydrogel remained almost constant throughout the tested frequency range (0.01–10 Hz) **(Figure S3B)**. Moreover, in comparison to the GelMA hydrogel, the CaNeu hydrogel could rapidly relax the external uniaxial compressive stress and retain only half of the residual elasticity, which indicates that the CaNeu hydrogel possesses considerably better network dynamics than the GelMA hydrogel **(Figure 1B)**. Furthermore, the CaNeu hydrogel exhibited significant shear-thinning properties with a “gel-sol” transition point at a shear strain of 300% **(Figure S3C)**. Owing to the good shear thinning and self-healing properties of the CaNeu hydrogel, it could be injected into a star-shaped mold using a 16G needle. The injected hydrogel remained free-standing in a star shape after the removal of the mold, and the majority of the encapsulated ADSCs remained viable after injection **(Figure 1C)**. The protection of ADSCs from the shear force during injection can be attributed to the reversible and alternating “gel-sol-gel” transition of the CaNeu hydrogel under conditions of oscillatory shear strain **(Figure 1D)**.

**Figure 1.**
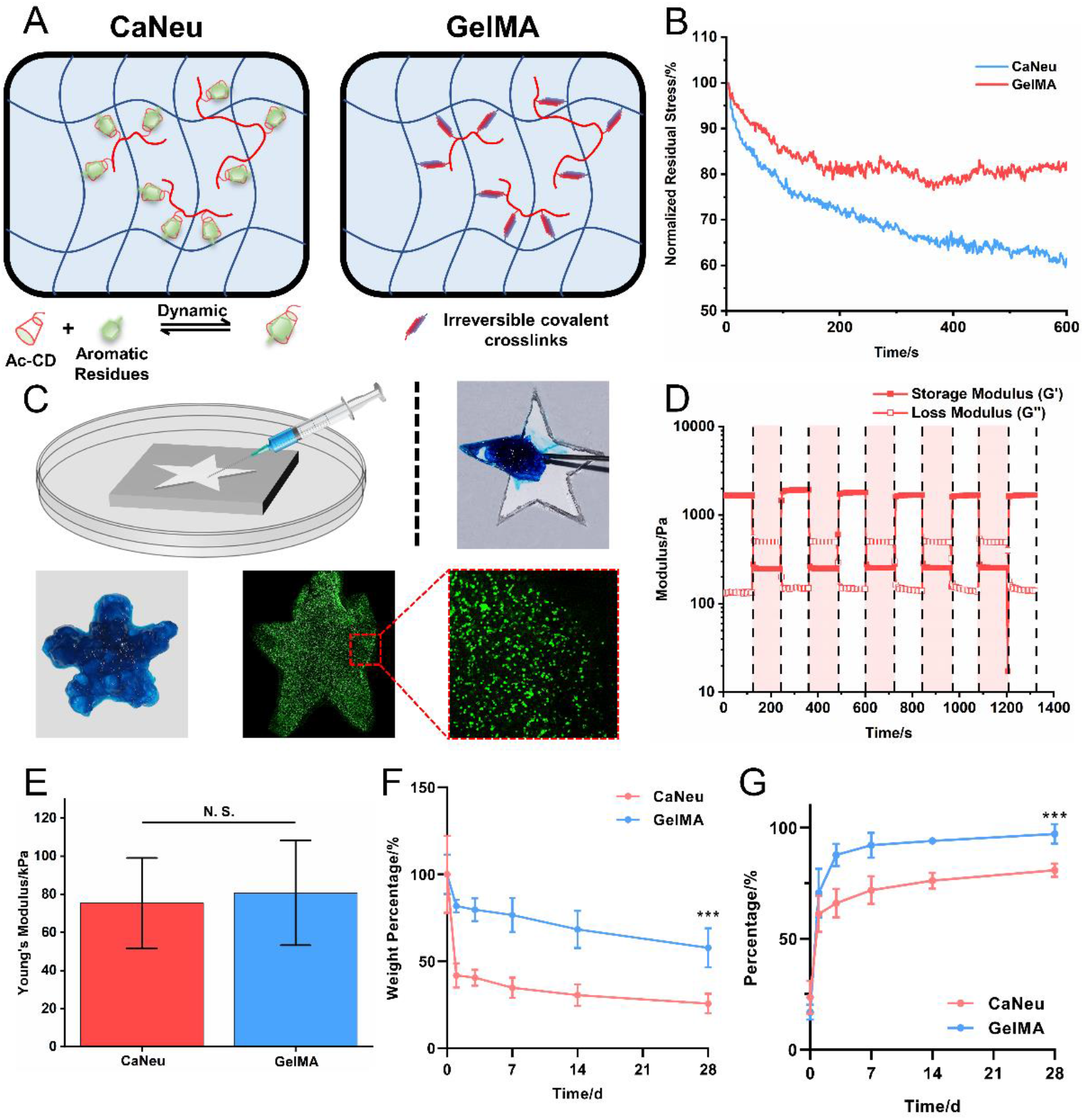
The CaNeu hydrogel exhibits highly dynamic behavior. (A) Schematic illustration of the structure of CaNeu and GelMA hydrogels. (B) The stress relaxation curves of CaNeu and GelMA hydrogels. The CaNeu hydrogel can relax higher compressive stress than the GelMA hydrogel within 15 min. (C) Demonstration of excellent injectability of the CaNeu hydrogel. Majority of the encapsulated ADSCs remained viable in the star-shaped mold after injection using a 16G needle. (D) The shear thinning properties of the CaNeu hydrogel. The CaNeu hydrogel showed fast and complete recovery under alternating high (500%) and low (1%) shear strain. (E) Young’s modulus of the CaNeu and GelMA hydrogels determined using a compression test. (F) The degradation profile of the CaNeu and GelMA hydrogels in PBS. (G) The release profiles of the encapsulated model protein (BSA) from the CaNeu and GelMA hydrogel (Student’s *t*-test; N. S.: no significance, \*\*\**P* < 0.001, data are shown as mean ± S. E. M.)

The CaNeu and GelMA hydrogels showed similar Young’s moduli without significant differences at the initiation of the degradation test **(Figure 1E)**. Micromorphological analysis revealed that the CaNeu and GelMA hydrogels exhibited similar network structures; however, the hydrogel network formed by the CaNeu hydrogels displayed more pores than the network formed by the GelMA hydrogels **(Figure S4)**. An appropriate degradation rate under physiological conditions is a prerequisite for the clinical application of the CaNeu hydrogels. The CaNeu hydrogel exhibited substantial degradation than the GelMA hydrogel and lost approximately 75% of its original weight after 21 days of degradation **(Figure 1F)**. However, the release of encapsulated bovine serum albumin (BSA) from the CaNeu hydrogel was slightly slower than that from the GelMA hydrogel, a phenomenon that might be attributed to the sequestration of proteins by the oligomerized CD nanoclusters **(Figure 1G)**[41]. These findings demonstrate the substantial dynamic behaviors of CaNeu hydrogels, including shear-thinning, self-healing, and injectability. We believe that the dynamic structure of the CaNeu hydrogel would provide a cell-adaptable microenvironment to support the growth and neurogenic differentiation of encapsulated ADSCs.

### 3.2 Dynamic CaNeu hydrogel supports ADSCs infiltration and viability

The cell migration assay **(Figure 2A)** revealed that ADSCs seeded on the surface of GelMA and CaNeu hydrogels remained largely on top of the former, while the majority of ADSCs infiltrated the latter **(Figure 2B, C)**. ADSCs viability was investigated by embedding single ADSCs within the two hydrogels, and then subjecting the cells to a live/dead cell viability assay **(Figure 2D)**. Live/dead cell staining revealed that the viability of ADSCs in the dynamic CaNeu hydrogel was higher than that in the GelMA hydrogel for 14 days **(Figure 2E)**. In addition, ADSCs in CaNeu exhibited a spread cell morphology, while those in GelMA remained round after 14 days of culture **(Figure 2F)**. These findings indicate that the ADSCs-loaded CaNeu hydrogels are cell-adaptable and support ADSCs penetration.

**Figure 2.**
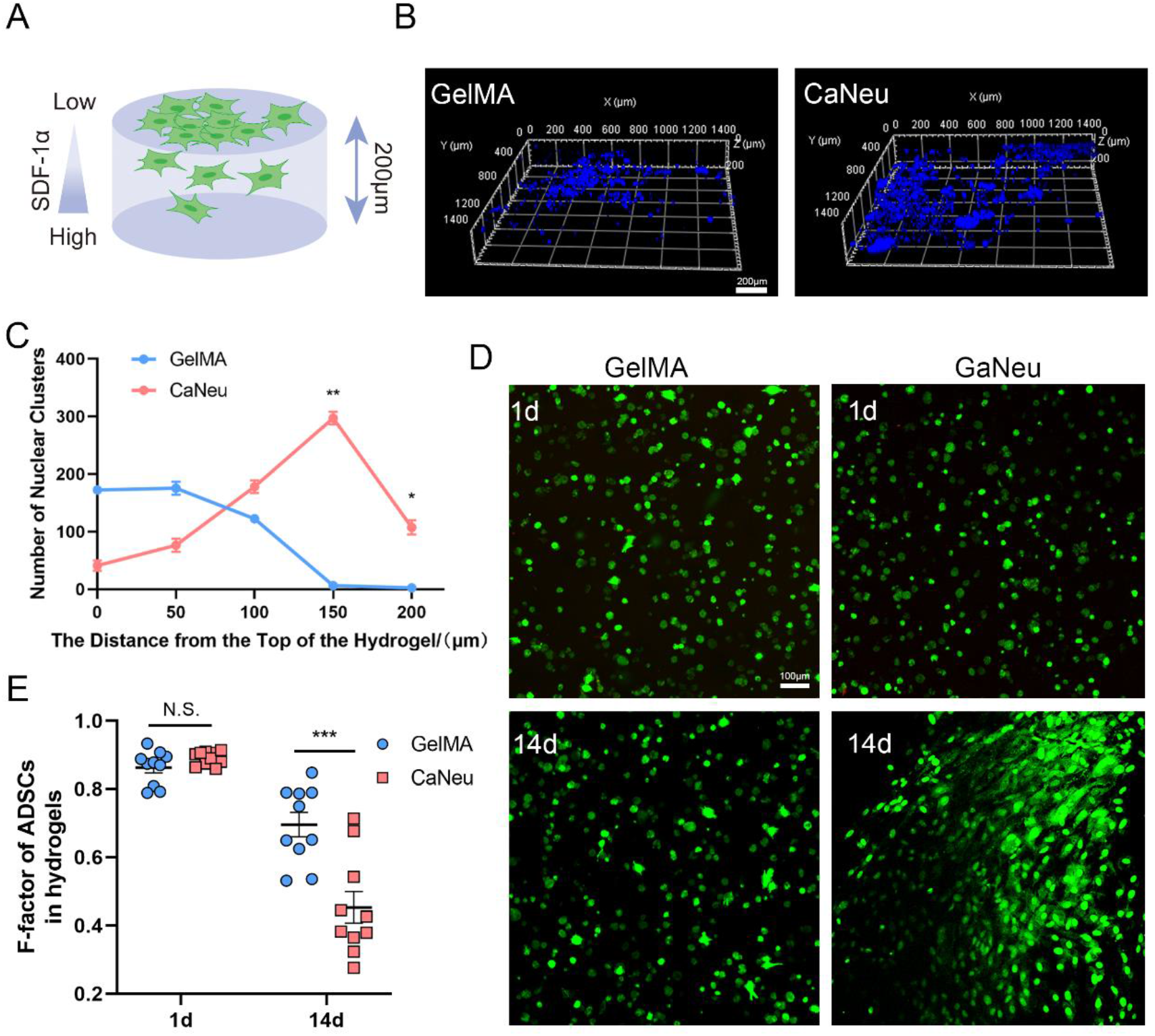
The cell-adaptable CaNeu hydrogel enables the rapid infiltration of external ADSCs and supports the viability of the encapsulated ADSCs. (A) Schematic illustration of the cell migration experiment conducted at 37℃. The ADSCs seeded on the upper surface of the CaNeu hydrogel infiltrated the hydrogels; however, only few seeded cells infiltrated the GelMA hydrogel. (B) Confocal micrographs and (C) quantification of the 3D distribution of DAPI-stained ADSCs nuclei within the GelMA and CaNeu hydrogels after 12 hours of exposure to a chemoattractant gradient at 37℃. Scale bar: 200 μm. (D) Viability of ADSCs in GelMA and CaNeu hydrogels after 1 and 14 days of *in vitro* culture. Scale bar: 100 μm. (E) Morphology of ADSCs was quantified using the F-factor (F = 4πA/P^2, where, A = area and P = perimeter). Smaller F-factor indicates greater degree of cell spreading (N. S.: no significance, \*\**P* < 0.01, \*\*\**P* < 0.001).

### 3.3 Dynamic CaNeu hydrogel enhances the stemness maintenance and neural differentiation of the encapsulated ADSCs^Sox2+^ through YAP-dependent signaling

Next, we examined the effects of the dynamic hydrogel structure on stemness maintenance and neural differentiation of encapsulated ADSCs^Sox2+^ in expansion and neuronal differentiation medium, respectively. The expression level of an ADSC^Sox2+^ stemness marker, nestin, sex-determining region Y-box transcription factor 2 (Sox2) **(Figure 3A, B)**and Vimentin **(Figure S5)**significantly increased in the CaNeu hydrogel with higher network dynamics after culturing ADSCs^Sox2+^ for seven days in expansion medium. Furthermore, the ADSCs^Sox2+^ in the CaNeu hydrogel exhibited substantially higher expression of the neuronal differentiation marker, microtubule-associated protein 2 (MAP2), than the ADSCs^Sox2+^ in the GelMA hydrogel after seven days of culture in neuronal differentiation medium **(Figure 3D)**. We also examined the expression of MAP2 and NeuN on one day, and found that the expression levels were low **(Figure S6)**. Moreover, owing to the enhanced neural differentiation, the ADSCs^Sox2+^ in the CaNeu hydrogel exhibited spread cell morphology, whereas the ADSCs^Sox2+^ in the GelMA hydrogel retained a round morphology **(Figure 3C, D)**.

**Figure 3.**
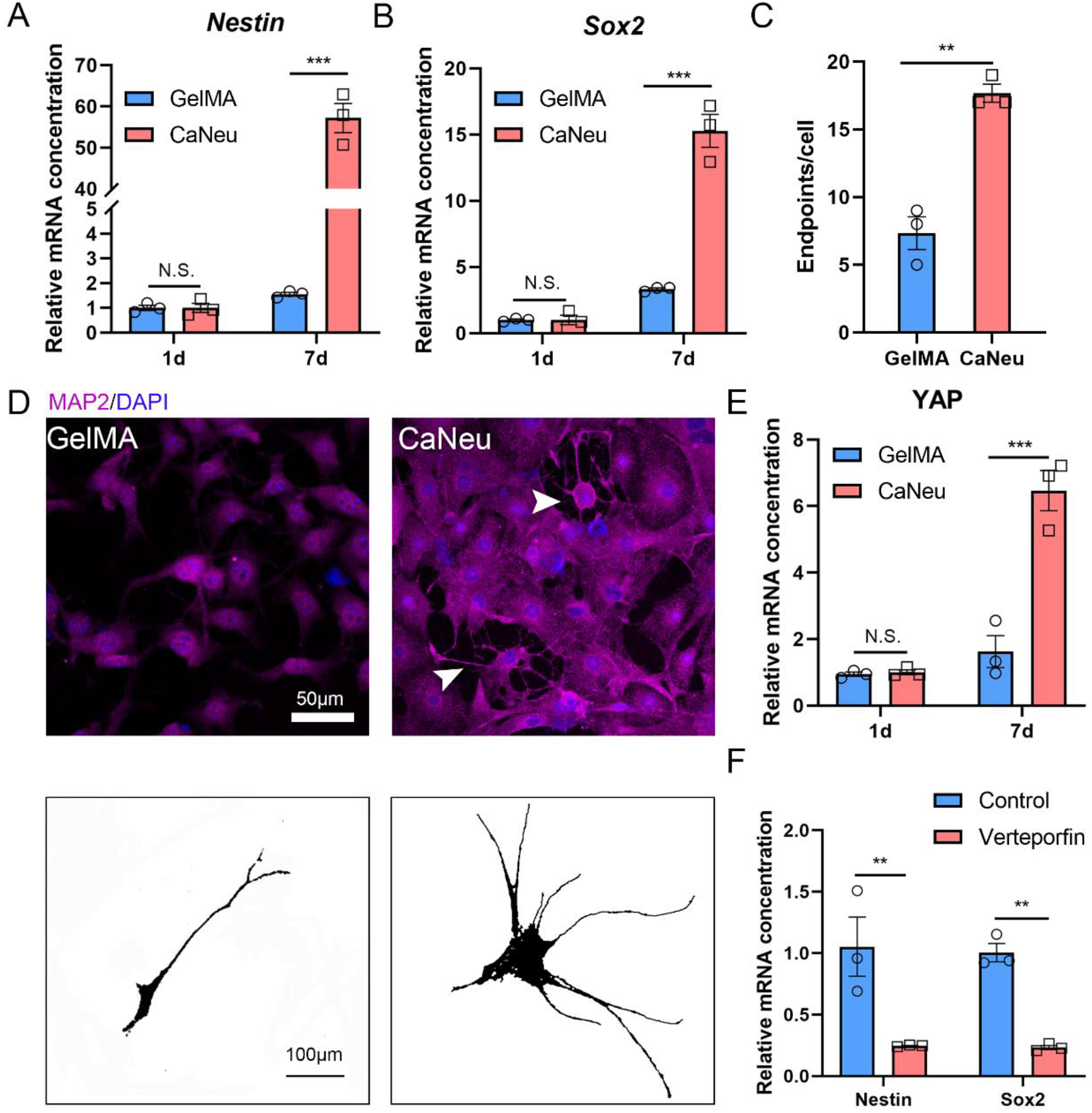
Cell-adaptable CaNeu hydrogel enhances neuronal differentiation and axonal growth of the encapsulated ADSCs^Sox2+^. (A, B) mRNA expression of stemness markers, nestin and Sox2, in ADSCs^Sox2+^ increased with the culturing time in the CaNeu hydrogel than that in the GelMA hydrogel (one-way ANOVA with Bonferroni’s post-hoc test; \*\**P* < 0.01, \*\*\**P* < 0.001, data are presented as mean ± S.E.M, n = 3). (C, D) Immunofluorescence to check for the expression of a neuronal differentiation marker, i.e., microtube-associated protein 2 (MAP2), in ADSCs^Sox2+^ encapsulated within the GelMA or CaNeu hydrogel and cell morphological reconstruction. Scale bar: 50 μm and 100 μm, respectively. (unpaired *t*-test, \*\**P* < 0.01, n = 3). (E) Expression of YAP mRNA in ADSCs encapsulated within the GelMA and CaNeu hydrogels (one-way ANOVA with Bonferroni’s post-hoc test; N. S.: no significance, \*\*\**P* < 0.001, n = 3). (F) Inhibition of YAP-TEAD interaction resulted in decreased expression of the stemness markers, nestin and Sox2 (one-way ANOVA with Bonferroni’s post-hoc test; \*\**P* < 0.01, data are presented as mean ± S.E.M, n = 3).

To study the mechanism underlying the maintenance of ADSCs^Sox2+^ stemness in the dynamic CaNeu hydrogel, we investigated Yes-associated protein (YAP)-dependent cell spreading. ADSCs^Sox2+^ showed markedly higher YAP expression in the CaNeu hydrogel after seven days of culture in the maintenance culture medium **(Figure 3E)**. Further, we blocked the YAP-TEAD (transcriptional enhanced associate domain) interactions by employing verteporfin, a small-molecule inhibitor, to investigate whether the maintenance of the stemness of the encapsulated stem cells involves YAP-dependent signaling. Blocking the YAP-TEAD interactions in the encapsulated ADSCs^Sox2+^ in the CaNeu hydrogel resulted in significant suppression of ADSCs stemness, as evidenced by the decreased expression of nestin, Sox2 and vimentin **(Figure 3F)**. These findings suggest that YAP-dependent signaling contributes to the enhanced stemness of the encapsulated ADSCs^Sox2+^ in the CaNeu hydrogel, which may result in the subsequent neural differentiation of ADSCs^Sox2+^ upon switching to the differentiation medium.

### 3.4 Delivery of ADSCs by CaNeu hydrogel promotes locomotor recovery in SCI rats

We further evaluated the efficacy of the ADSCs-loaded CaNeu hydrogel (CaNeu/ADSCs) at repairing SCI in a rat model **(Figure 4A)**. In comparison to the hindlimbs that were incapable of supporting weight in the SCI group, the animals in the CaNeu/ADSCs group exhibited higher support of weight by the hindlimb and occasional coordination of the hindlimb and forelimb **(Figure 4B, Supplementary video)**. Functional recovery of hindlimb locomotion was assessed using the Basso, Beattie, and Bresnahan (BBB) scoring system eight weeks after surgery **(Figure 4C)**. The animals in the CaNeu/ADSCs group exhibited significant functional recovery than those in other groups. Functional score of the CaNeu/ADSCs group reached a mean value of 9.2 ± 2.0 on the BBB scale, which is markedly higher than that of the cell-free CaNeu hydrogel-treated (5.0 ± 2.8) and ADSCs-treated groups (4.5 ± 3.2) **(Figure 4C)**. Moreover, electrophysiological analysis was performed to evaluate neural relay restoration at the lesion site after eight weeks **(Figure 4D)**. Compared with the baseline response in the SCI group, rats in the CaNeu/ADSCs group achieved a mean motor evoked potential (MEP) amplitude of 0.11 ± 0.04 mV, which was higher than that of cell-free CaNeu hydrogel-treated (0.07 ± 0.02 mV) and ADSCs (0.06 ± 0.02 mV) groups, indicating enhanced recovery of evoked responses by ADSCs-loaded hydrogels **(Figure 4E, F)**. Re-transection of the spinal cord at the T10 level (rostral to the implant site) resulted in the loss of evoked potentials in the hindlimbs **(Figure S7B)**, confirming the formation of new electrophysiological relays across the lesion center.

**Figure 4.**
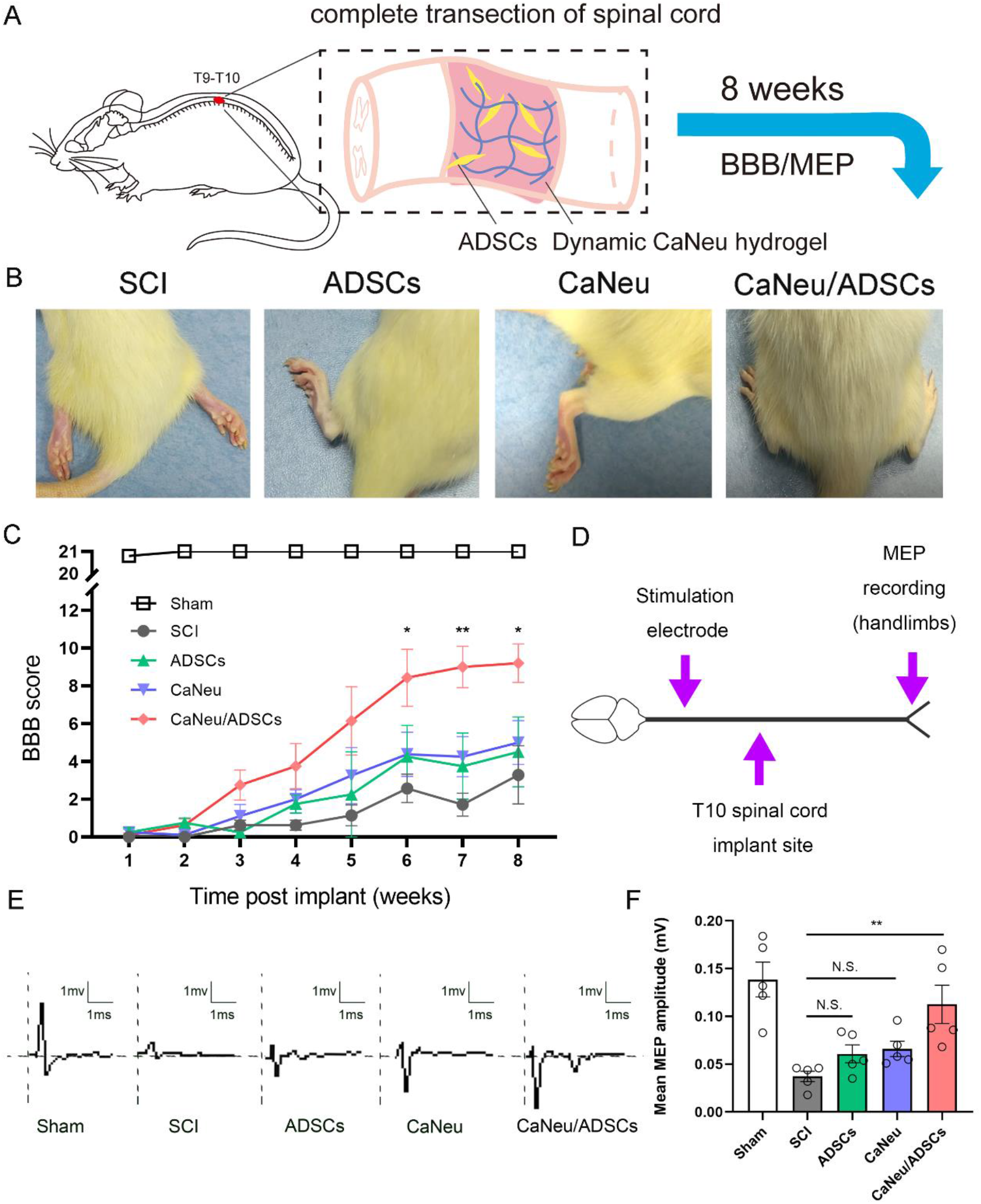
Transplantation with ADSCs-loaded CaNeu hydrogel promotes electrophysiological and locomotor recovery in SCI rats. (A) Schematic illustration of the workflow of the therapeutic experiment. (B) Representative records of animal walking gaits eight weeks post-SCI revealed differential hindlimb walking patterns of the sham control, SCI, ADSCs-treated, CaNeu-treated, and ADSCs-loaded hydrogel-treated (CaNeu/ADSCs) groups. (C) Comparison of locomotor recovery for the SCI, ADSCs, CaNeu, and CaNeu/ADSCs groups. Locomotor recovery was measured using the BBB scale in an open field (Tukey’s multiple comparison test in two-way ANOVA; \**P* < 0.05, \*\**P* < 0.01; mean ± S.E.M, n = 9 animals). (D) Schematic diagram depicting the workflow of the electrophysiology study performed at eight weeks post implantation. The rostral extremity of the spinal cord region was exposed to stimulation electrode, and motor evoked potentials (MEPs) were recorded in the hindlimbs. (E) Rats in the CaNeu/ADSCs group exhibited evident MEP responses, while those in the SCI group showed MEP at the baseline level. (F) The mean MEP amplitude of animals in CaNeu/ADSCs group was significantly higher than that of animals in other groups (one-way ANOVA followed by Tukey’s post hoc analysis; N. S. = no significance, \*\**P* < 0.01, mean ± S.E.M, n = 5 animals).

### 3.5 Delivery of ADSCs via the CaNeu hydrogel facilitates neurogenesis and axonal regeneration in SCI rats

We performed hematoxylin and eosin (H&E) staining and immunostaining to assess the native tissue response, including the volume of the cystic cavities, eight weeks after SCI **(Figure S8A)**. The mean volume of cystic cavities was significantly reduced in the CaNeu/ADSCs group compared to SCI group (0.28 vs. 0.02 mm^2^, *P* < 0.001) **(Figure 5A, D)**. Moreover, the cell-free CaNeu hydrogel-treated group exhibited a faster degradation rate *in vivo* than the CaNeu/ADSCs) group **(Figure S8A)**. The difference degradation rate due to the cell-laden hydrogel contains the cell-secreted extracellular matrix molecules which may help stabilize the hydrogel and slow down the degradation [42, 43]. In the CaNeu/ADSCs group, the central lesion was filled with an extracellular matrix (ECM)-like structure, which was mostly devoid of astrocytes but was surrounded by astrocytic scars, as revealed by H&E staining and glial fibrillary acidic protein (GFAP, an astrocyte marker) immunostaining **(Figure 5A**, **Figure S8A**). Immunostaining revealed that CaNeu/ADSCs transplantation resulted in a considerable increase in the proportion of TUBB3^+^ structures in the central lesion (47.64% ± 3.65%) compared to that in the CaNeu-treated (43.59% ± 1.27%), ADSCs-treated (32.80% ± 1.41%), and SCI groups (18.21% ± 1.86%) **(Figure 5E)**. A significantly larger number of myelin basic protein (MBP; a myelin marker) positive cells was observed at the lesion epicenter in the CaNeu/ADSCs group **(Figure S8B, C)**. Further, we stained Neurofilament 200 (NF200, an axonal marker) to identify mature neurons with functional axons eight weeks after injury. NF200 staining showed the regenerated axons in the rostral border and lesion center in the CaNeu/ADSCs group **(Figure S9A, B)**. Host serotonergic axons also regenerated into the hydrogel **(Figure 5B)**, and the number of serotonergic axons (45 ± 5) labeled with 5HT observed at the lesion and boundary of the spinal cord sections in the CaNeu/ADSCs group was significantly higher than that in the ADSCs-treated (15 ± 3) and CaNeu-treated groups (18 ± 1) **(Figure 5F)**. Additionally, transmission electron microscopy revealed regeneration of the myelin sheath in the damaged area **(Figure 5C)**and improved structural integrity of axons in the CaNeu/ADSCs group than that in the SCI group. These results showed that the CaNeu hydrogel in combination with ADSCs promoted neurogenesis and axonal regeneration in an animal model of SCI.

**Figure 5.**
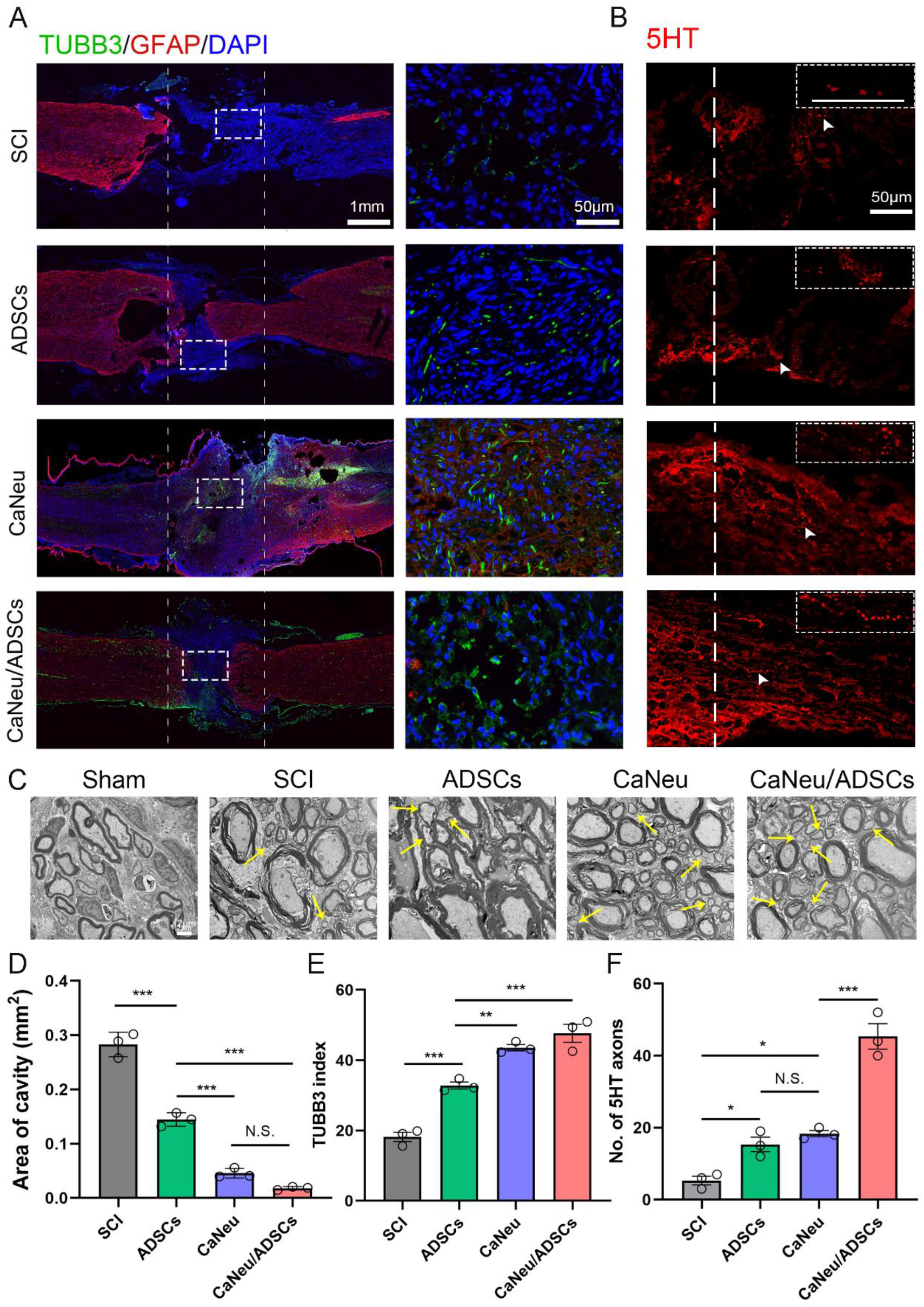
CaNeu-hydrogel‒mediated ADSCs delivery reduces the volume of cavities and enhances axonal regrowth and remyelination in SCI rats. (A) Representative confocal immunofluorescence images of the spinal cord sections revealing the distribution of TUBB3^+^ structures (TUBB3, green) and astrocytes (GFAP, red) in all groups 60 days post-injury. Scale bar: 1 mm and 50 μm, respectively. (B) Representative images of immunostaining for 5HT, an inhibitory neurotransmitter released by serotonergic axons, in spinal cord sections of the lesion epicenter from animals of all groups. Scale bar: 20 μm. (C) Electron microscopy images revealed that transplantation of CaNeu loaded with ADSCs induced remyelination after SCI. The yellow arrows indicate regenerated myelin whose myelin lamellae were thin but tight. Scale bar: 2 μm. (D) Graphs showing the quantified volumes of cavity. (E) Quantification of TUBB3 positive staining per mm2 of tissue in the spinal cord sections (one-way ANOVA followed by Tukey’s post hoc analysis; N. S.: no significance, \*\**P* < 0.01, \*\*\**P* < 0.001, error bars represent the S.E.M, n = 3 animals per group). (F) Quantification of 5-HT positive neurons at the lesion site (one-way ANOVA analysis; N. S.: no significance, \*\**P* < 0.01, \*\*\**P* < 0.001, error bars represent the S.E.M, n = 3 animals per group).

### 3.6 Delivery of ADSCs via the CaNeu hydrogel results in the suppression of apoptosis and inflammation in SCI through the promotion of M2 polarization of resident macrophages via inhibition of the PI3K/Akt signaling pathway

It has been reported that soft implant materials reduce expression of inflammatory markers in surrounding macrophages when compared to stiff materials [44] and ADSCs may play an important role in immunomodulation by secreting paracrine signals [45]. Therefore, we hypothesize that CaNeu/ADSCs treatment promotes recovery after SCI by suppressing apoptosis and inflammation at the lesion site. Immunostaining for cleaved caspase-3 (C-Caspase3) at seven days after injury revealed that CaNeu/ADSCs treatment resulted in the presence of minimal number of C-Caspase3^+^ cells at the site of the injury **(Figure 6A, C)**. Moreover, western blotting revealed the expression of Bax, which could promote apoptosis, was significantly attenuated by CaNeu/ADSCs treatment **(Figure 6D, E)**. Meanwhile, the CaNeu/ADSCs-treated group exhibited an increase in the level of anti-apoptosis protein Bcl-2 compared with that in the SCI group **(Figure 6F)**. In addition, the expression of IBA1, a marker of inflammatory macrophages and microglia, was used to assess the reactive inflammatory responses. One week after SCI, the intensity of IBA1 immunoreactivity was significantly attenuated at the border surrounding the newly formed tissue in the central lesion of the CaNeu/ADSCs group **(Figure S10 A, B)**.

**Figure 6.**
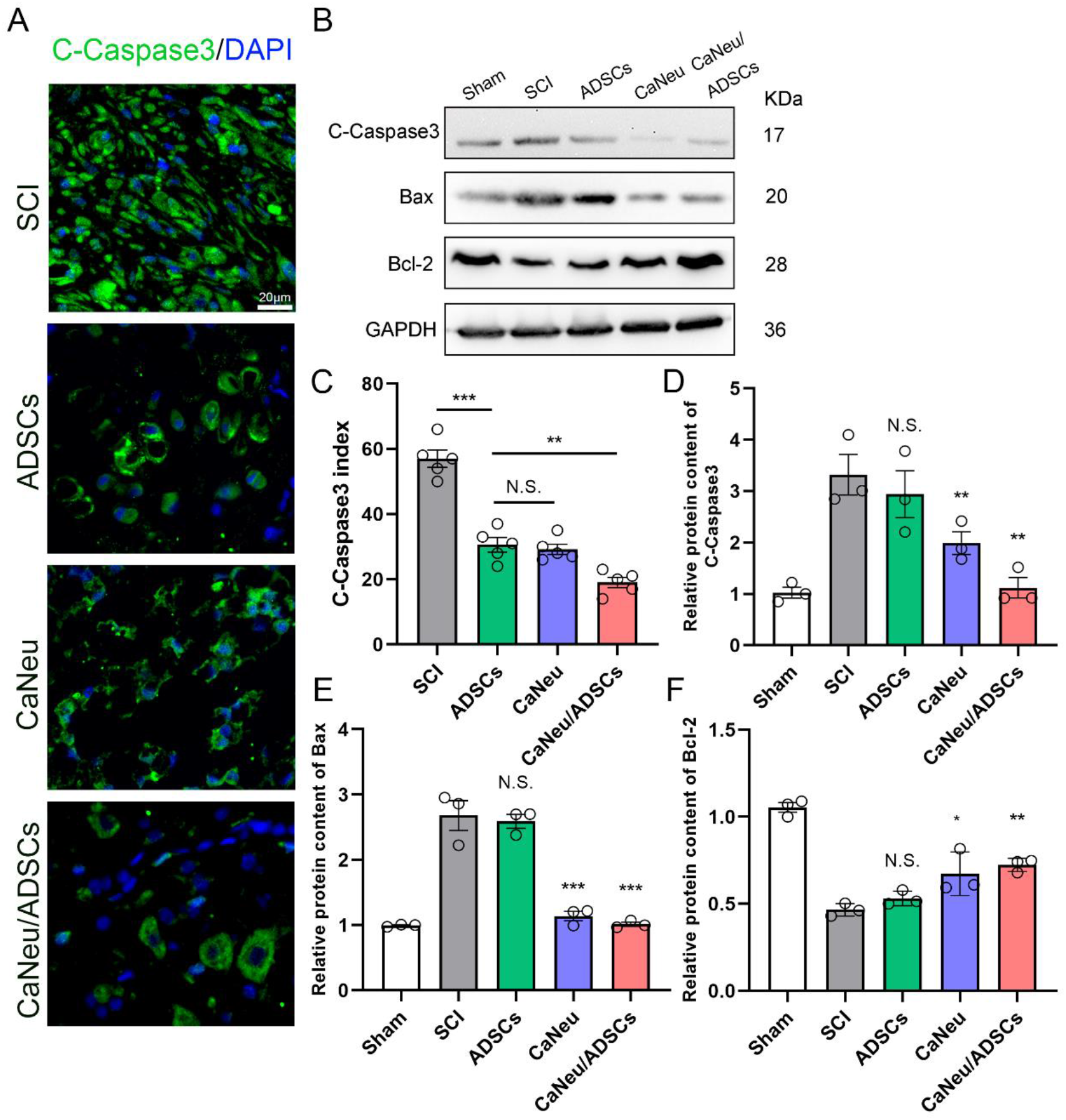
Transplantation with ADSCs-loaded CaNeu hydrogel alleviates apoptosis in a rat SCI model. (A) Confocal immunofluorescence images showing the expression of the apoptotic marker cleaved caspase-3 seven days post-injury (one-way ANOVA followed by Tukey’s post hoc analysis; \**P* < 0.05, \*\**P* < 0.01, error bars represent S.E.M, n = 5 animals per group). Scale bar: 20 μm. (B, D, F) Western blots and quantification data of cleaved caspase-3, Bax, and Bcl-2 in each group seven days after surgery. (C) Quantification of caspase-3 immunofluorescence one week after injection (one-way ANOVA followed by Tukey’s post hoc analysis; N. S.: no significance, \**P* < 0.05, ** *P* < 0.01, error bars represent the S.E.M., n = 3 animals per group).

Next, we examined macrophages at the lesion epicenter, and observed considerable accumulation of CD68^+^ cells in the CaNeu/ADSCs group seven days after SCI **(Figure 7A)**. Interestingly, in the first week post-injury, CD163 was only expressed in the CaNeu/ADSCs group, while in the second week, the expression of CD163 was observed in all groups **(Figure 7A)**. Therefore, we performed double staining of CD68 and CD163, and observed that the CaNeu/ADSCs group was characterized by the presence of a large population of M2 macrophages at the lesion site **(Figure 7A, B)**. To further elucidate the mechanism underlying the anti-inflammatory effect of the CaNeu/ADSCs treatment, we explored the potential involvement of the PI3K/Akt pathway. Expression of pro-inflammatory cytokines IL-6 and CD206 (a marker of M2 macrophages) was examined using western blotting. The SCI group exhibited increased expression of IL-6, which was sharply reduced in response to CaNeu/ADSCs treatment **(Figure 7C, D)**. In addition, higher expression of CD206 and reduction of pAkt was observed in the CaNeu/ADSCs group than that in the SCI group **(Figure 7C-F)**, suggesting that the therapeutic effect of CaNeu/ADSCs is mediated via the IL-6/PI3K/Akt signaling pathway. Collectively, these data suggest that the dynamic CaNeu hydrogel in combination with ADSCs alleviates neuroinflammation through the PI3K/Akt pathway to promote axonal regeneration, myelination, and locomotor function recovery.

**Figure 7.**
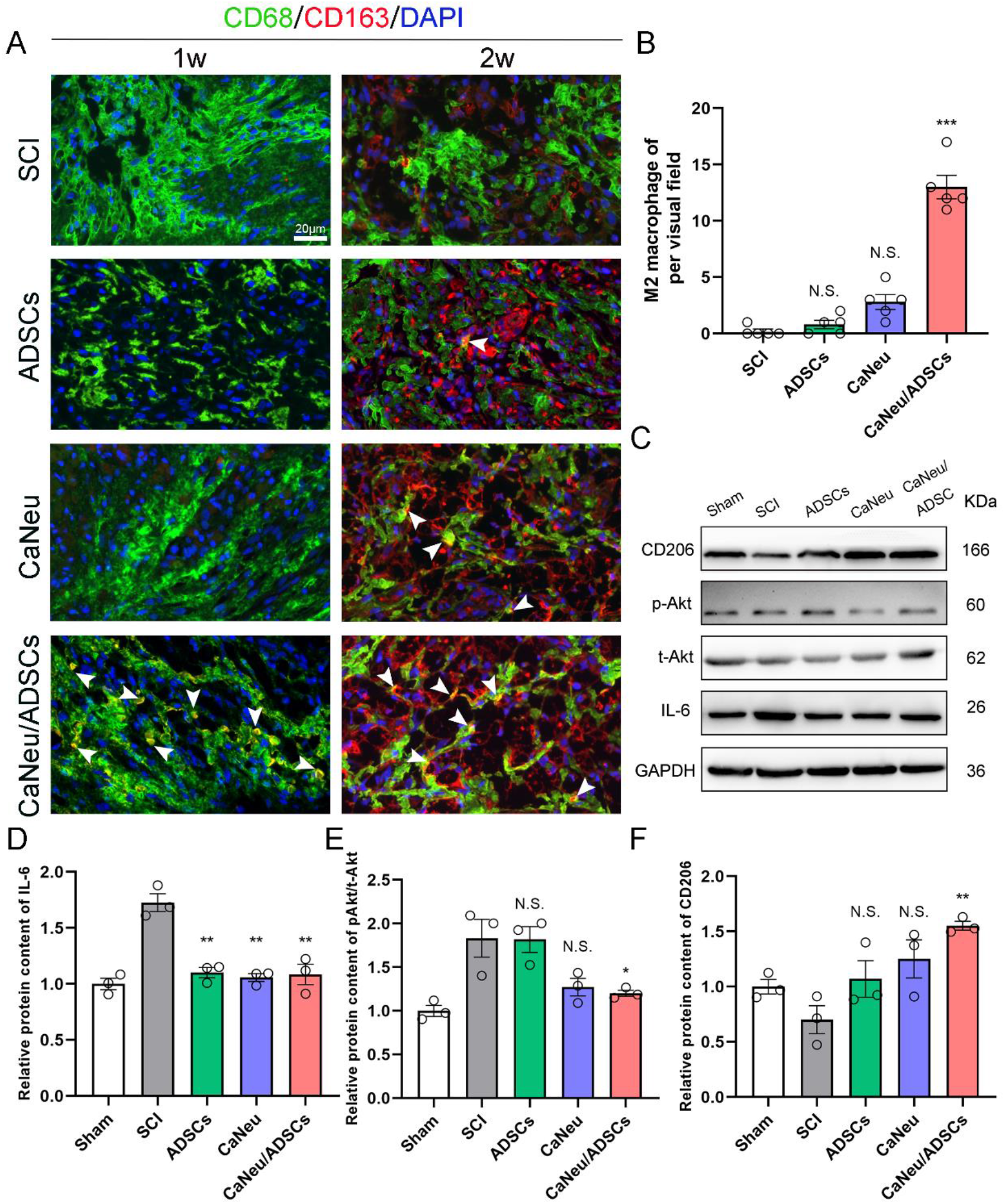
Transplantation with ADSCs-loaded CaNeu hydrogels promotes M2 polarization of the recruited macrophages by activating the PI3K/Akt pathway after acute traumatic SCI rats. (A) Confocal immunofluorescence microscopy images showing the increased presence of M2 macrophages in the CaNeu/ADSCs group, as evidenced by the co-staining of CD68 and CD163 (arrows represent M2 macrophages). Scale bar: 20 μm. Data are expressed as mean ± S.E.M. (B) Percentage of M2 macrophages two weeks after SCI (one-way ANOVA followed by Tukey’s post hoc analysis; N. S.: indicates no significance, \*\*\**P* < 0.001, error bars represent the S.E.M., n = 5 replicate experiments per group). (C–F) Western blots and quantification data for IL-6, pAkt, and CD206 (a marker of M2 macrophages) in each group one week after SCI (one-way ANOVA followed by Tukey’s post hoc analysis; N. S.: no significance, \**P* < 0. 05, \*\**P* < 0. 01, data represents the mean ± S.E.M, n = 3).

## 4. Discussion

Acute SCI is increasingly being recognized as the cause of severe and permanent disability. Stem-cell‒based therapy is an attractive strategy for spinal cord regeneration and functional recovery. Zhou *et al.* [46] reported that ADSCs transplantation alleviates SCI-induced neuroinflammation to promote SCI repair; however, experimental results reveal the persistence of a cavity even after ADSCs transplantation. The limited efficacy of direct stem cell transplantation may be attributed to the low retention and survival rates of the delivered stem cells. Therefore, a combination of stem cells and supportive biomaterials might enhance the therapeutic outcomes of stem cell transplantation.

Cells in living tissues continuously interact with neighboring cells and the ECM to perform various activities, including spreading, migration, proliferation, and differentiation [47]. Hydrogels that recapitulate the key features of natural ECM represent attractive candidates to be used in tissue engineering and regenerative medicine [48]. Therefore, the design of dynamic hydrogels mimicking the viscoelastic properties of natural ECM is crucial for direct expansion and differentiation of encapsulated stem cells, thereby facilitating the regeneration and functional recovery of damaged spinal cord tissues. Compared with that of the covalent GelMA hydrogel, the network formed by the developed CaNeu hydrogel contains dynamic host–guest crosslinks, which can be disassociated and re-associated in a reversible manner. The rapid stress relaxation can be attributed to the highly dynamic hydrogel network, where the applied external forces are attenuated through chain deformation and rearrangement. The traction forces exerted by the encapsulated cells can facilitate such hydrogel network reorganization, thereby generating the space required for cell growth and spreading [42, 49, 50]. Furthermore, the neural differentiation of the ADSCs encapsulated in the CaNeu hydrogel requires the dynamic hydrogel network to support cell spreading and extension of axonal protrusions. Therefore, the dynamic CaNeu hydrogel with rapid stress relaxation can effectively support the neural differentiation of the encapsulated ADSCs. We found that ADSCs^Sox2+^ expressed elevated levels of YAP when encapsulated within the dynamic CaNeu hydrogel, indicating the enhanced mechanosensing of ADSCs [51]. In addition, the transcriptional co-activator YAP has been reported to be involved in determining the survival, expansion, and differentiation of multiple stem cell types [52]. Previous studies have revealed that appropriate expression of YAP plays an important role in the maintenance of stemness and differentiation of embryonic and induced pluripotent stem cells [53, 54]. Blocking of YAP-TEAD interactions revealed that maintenance of stemness and differentiation of ADSCs^Sox2+^ relied on YAP-dependent signaling. This finding reveals the ability of the CaNeu hydrogel to facilitate stem cell differentiation.

Acute SCI can induce the generation of a complex neuroinflammatory microenvironment, resulting in impaired recovery from SCI and progressive tissue degeneration [55, 56]. Previous studies have shown that M2 macrophages are essential for tissue healing and regeneration [57–59]. We found that the dynamic CaNeu hydrogel loaded with ADSCs significantly increased the population of M2 macrophages, indicating that this might be the mechanism by which combinational transplantation exerts its effects. Moreover, the co-administration of CaNeu hydrogel and ADSCs can minimize the deleterious effects of inflammation at an early stage and promote nerve fiber regeneration. Previous studies have demonstrated that biophysical cues could modulate phenotype selection of the recruited macrophages in the hydrogels [60, 61]. Meli *et al.* [44] reported that a soft hydrogel matrix enhanced adhesion and suppressed the M1 polarization of macrophages. Previous studies have also demonstrated the immunomodulatory potential of ADSCs via paracrine signals, including growth factors, cytokines, and exosomes [45]. Kruger *et al.* [62] reported that ADSCs-conditioned media elicited an anti-inflammatory immune response. Therefore, we speculated that the dynamic CaNeu hydrogel network provided an immunosuppressive microenvironment for the recruited host macrophages. The reduced M1 population secreted substantially less IL-6, which further consolidated the anti-inflammatory microenvironment. Previous studies have suggested that IL-6, a pro-inflammatory factor, may play an important role in the phenotypic selection of the recruited macrophages in SCI [63–65]. Therefore, we investigated the downstream signaling pathways activated by IL-6. We found that the PI3K/Akt signaling pathway was activated when the expression of IL-6 was significantly inhibited in the CaNeu hydrogel loaded with ADSCs. Nevertheless, our study has certain limitations, for example, the lack of tracking neuronal differentiation of transplanted ADSCs *in vivo*. In subsequent studies, we aim to label ADSCs with a nanoprobe [66] to further improve the regenerative effects of co-administration of CaNeu hydrogel and ADSCs. This may potentially assist in exploring reliable, efficient, and accurate strategies for neuron regeneration and functional recovery after SCI.

## 5. Conclusion

In summary, we designed a cell-adaptable CaNeu hydrogel as a vehicle for the delivery of ADSCs. The dynamic network of the CaNeu hydrogel provides a favorable microenvironment for enhanced spreading of encapsulated ADSCs via YAP-dependent signaling. More importantly, paracrine factors secreted by ADSCs triggered PI3K/Akt pathway and enhanced polarization of M2 of recruited macrophages, thereby significantly alleviate neuroinflammation at the site of injury. Our strategy of employing a dynamic CaNeu hydrogel loaded with ADSCs can potentially enable stem-cell-based therapy for acute SCI, and opens new avenues for dynamic hydrogel platforms aimed at the treatment of other central nervous system diseases and injuries.

## Acknowledgement

This work was supported by grants from National Natural Science Foundation of China (Grant No.82071362). This work is also supported by the Basic Research Project Fund, the Government of the Shenzhen Science and Technology Innovation Commission (Project No. JCYJ20190809165201646). The authors would like to thank Jing Zhao, Krsna for revising the manuscript and providing valuable suggestion.

## Supporting Information

Supplementary data includes ten Figures and one table.

