## Supplemental Figures for "Cell-adaptable dynamic hydrogel reinforced with stem cells improves the functional repair of spinal cord injury by alleviating neuroinflammation": biorxiv-supplementary files.docx

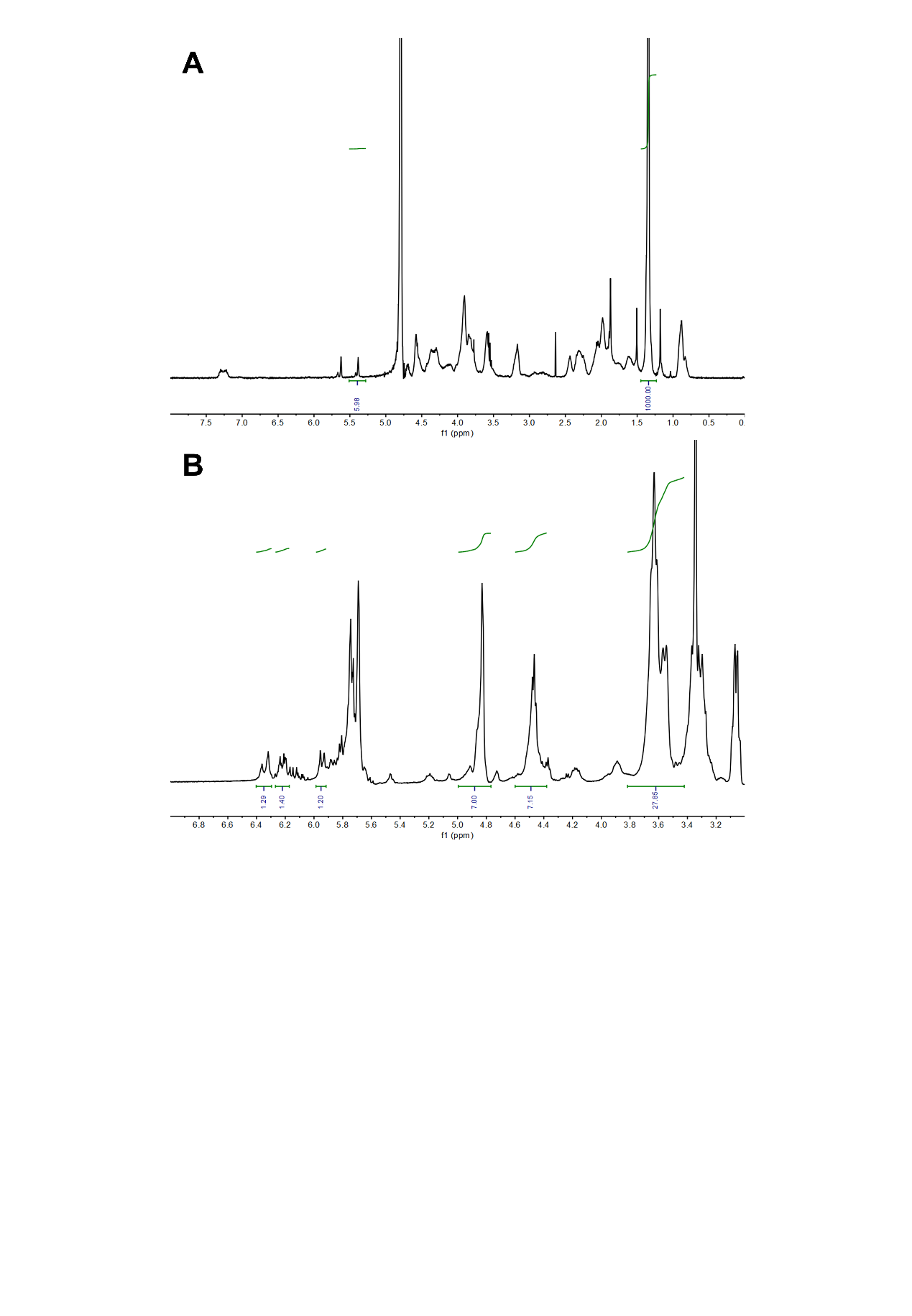

**Figure S1. The NMR analysis.** (A) GelMA, (B) Ac-β-CD.

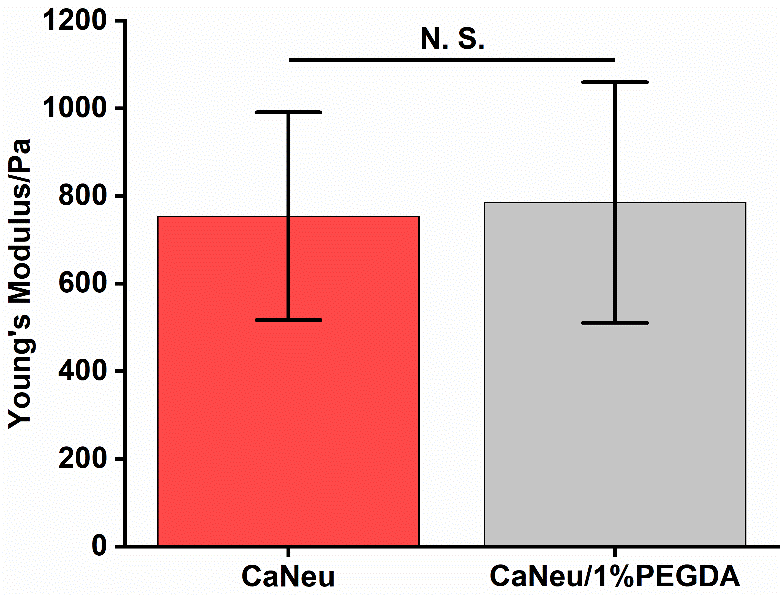

**Figure S2. The compression tests of CaNeu hydrogel and CaNeu/1% PEGDA hydrogel**

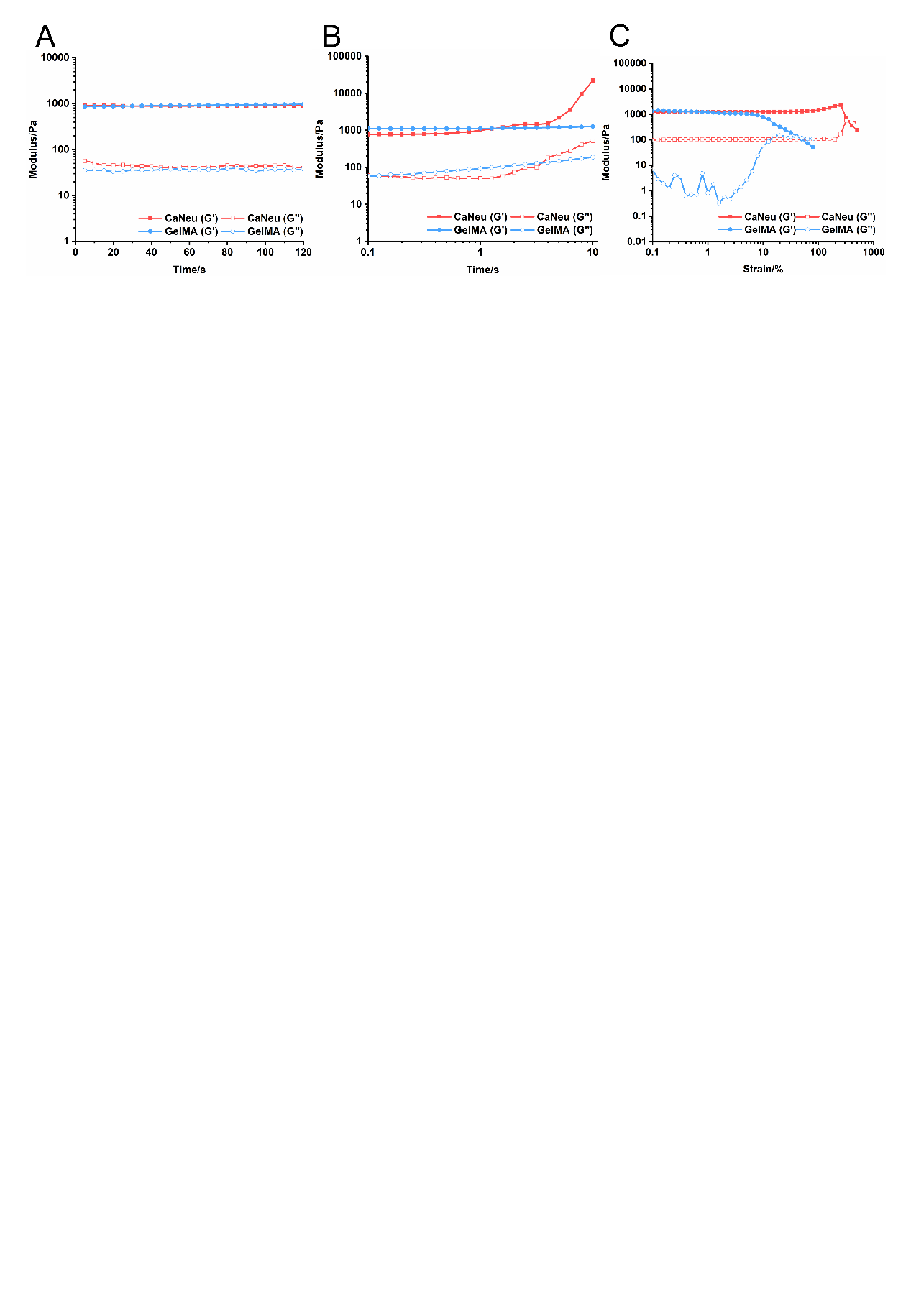

**Figure S3. The Rheological test of CaNeu and GelMA hydrogel.** (A) The timey sweep of CaNeu and GelMA hydrogel. (B) The frequency sweep of CaNeu and GelMA hydrogels from 0.01 Hz to 10 Hz. (C) The strain sweep of CaNeu and GelMA hydrogels. The CaNeu hydrogels showed significant higher tenacity as revealed by the much higher elongation at break without comprising their stiffness.

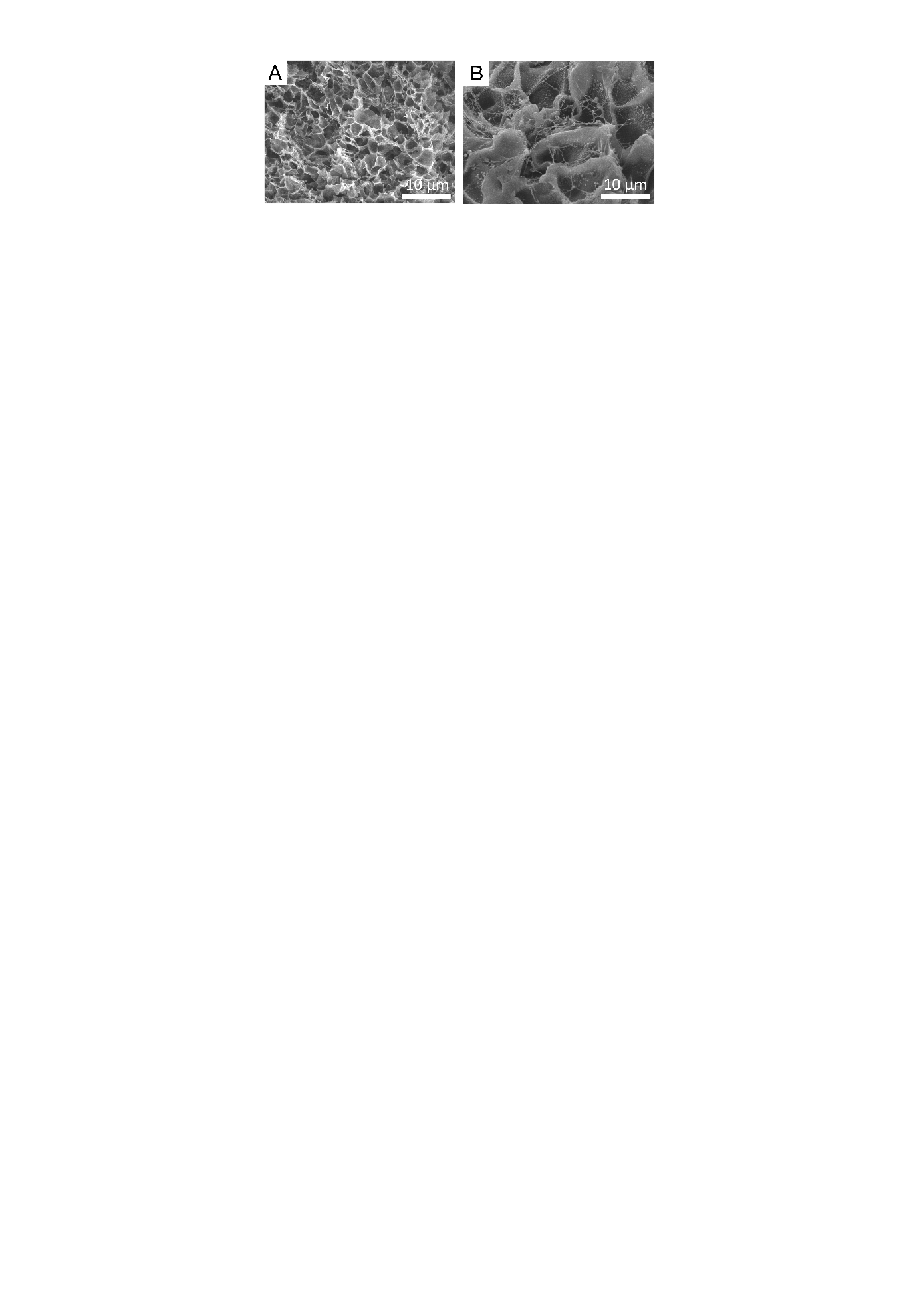

**Figure S4. The SEM images of** (A) CaNeu hydrogel and (B) GelMA hydrogel

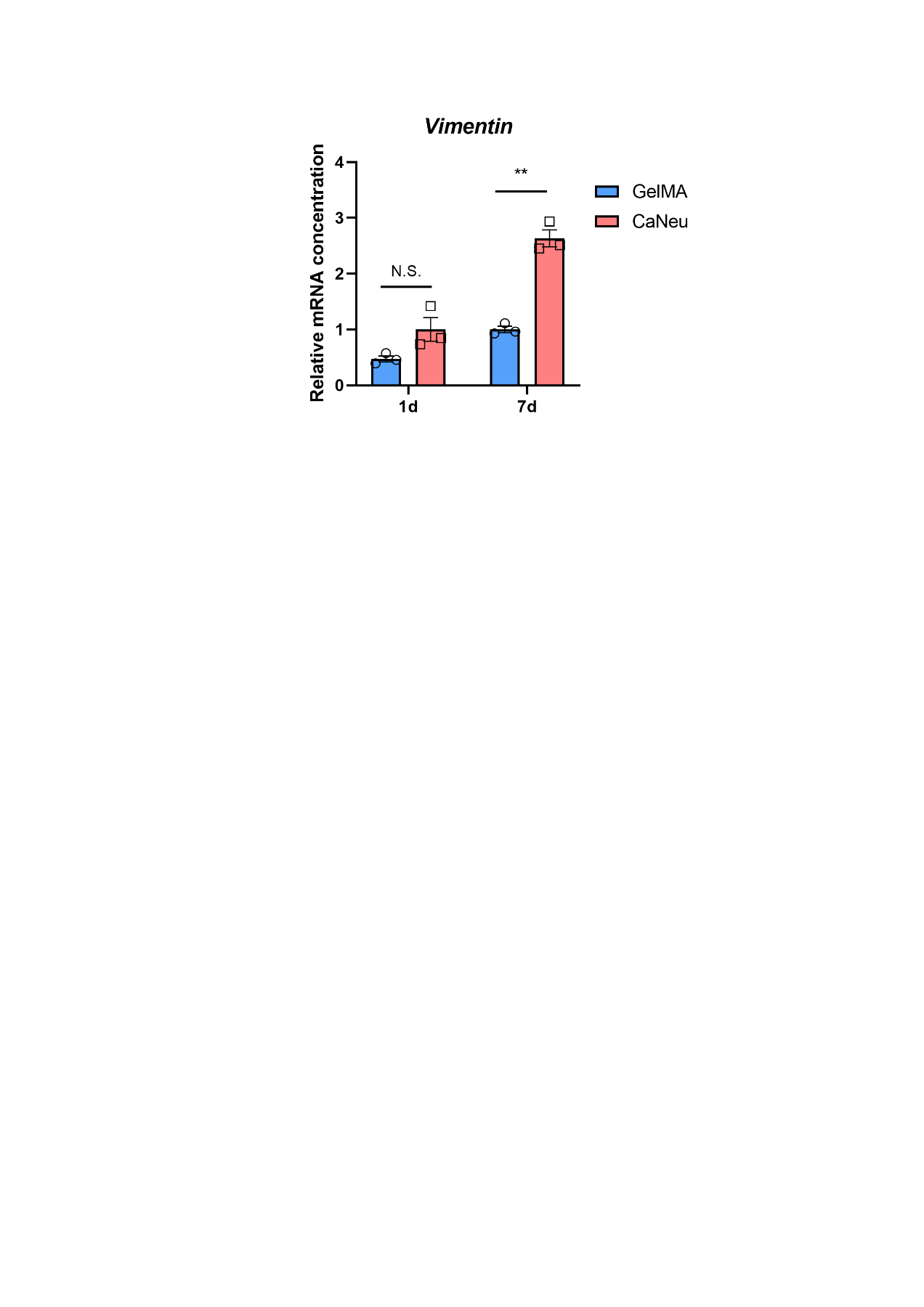

**Figure S5.** Quantitative gene expression levels of the *Vimentin* as determined by qPCR in the stem cells encapsulated in GelMA and CaNeu hydrogels after 1d and 7d in expansion medium.

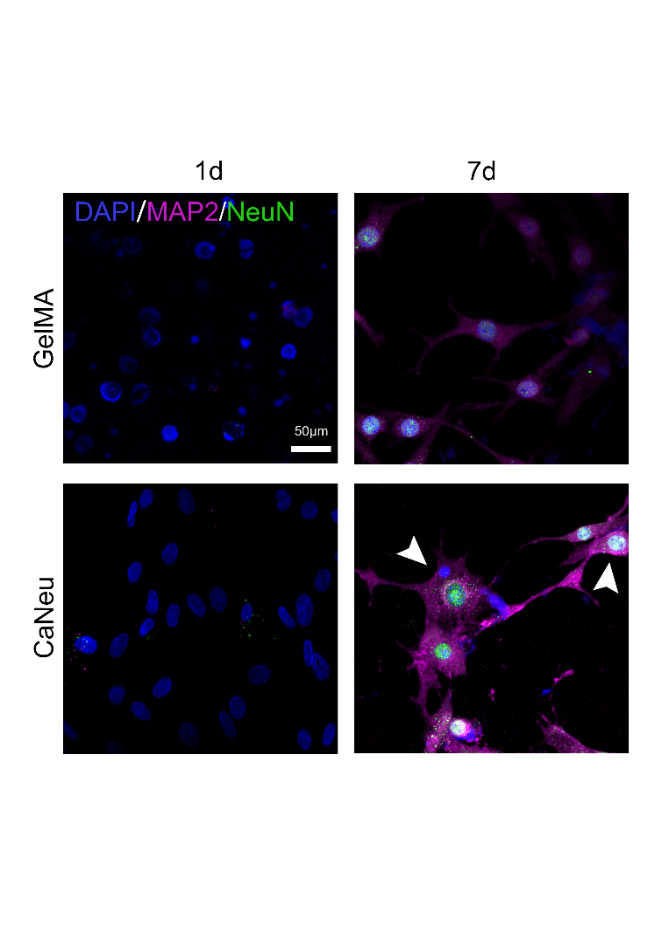

**Figure S6.** **CaNeu hydrogel enhances neuronal differentiation of the encapsulated cells.** Only after 7d by induction of neuronal differentiation, do encapsulated cells stain positive for the differentiated neuronal markers microtubule-associated protein 2 (MAP2) and NeuN, while on the 1d there were few MAP2^+^/NeuN^+^ cells in both CaNeu and GelMA groups. Scale bar, 50μm.

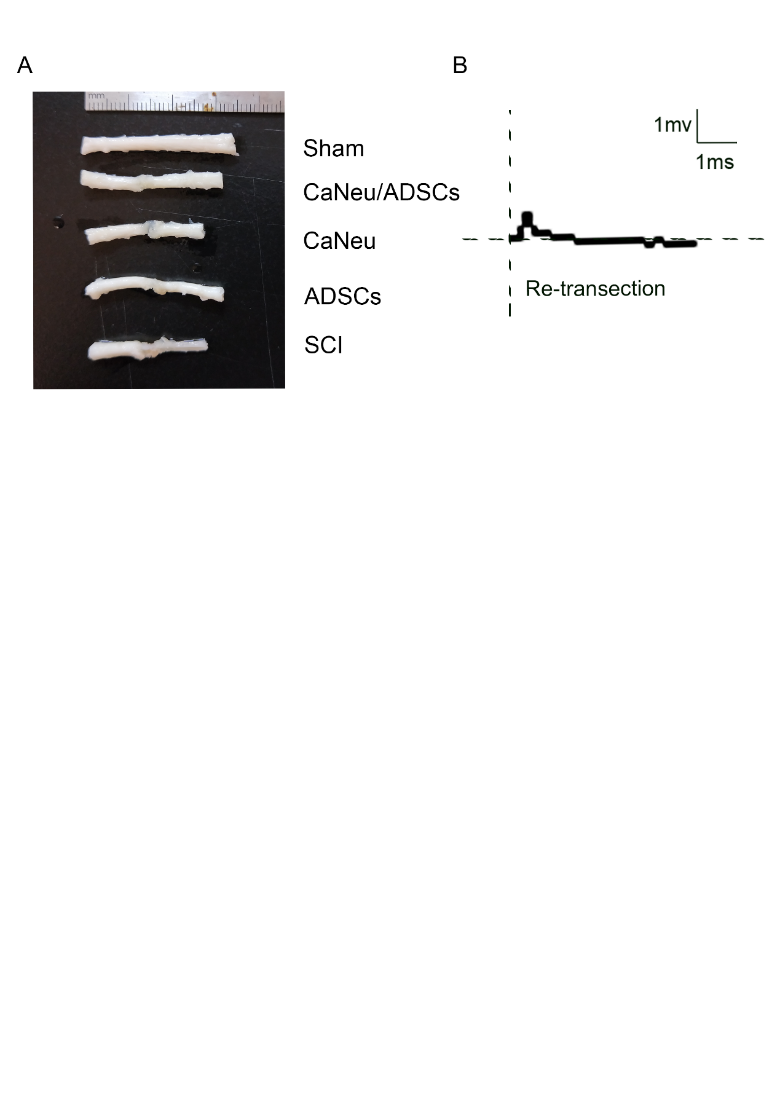

**Figure S7. Gross specimen and electrophysiological re-transection of rat spinal cord.** (A) Gross image of the complete transection spinal cord of rats, eight weeks post-injury. (B) CaNeu loaded with ADSCs exhibit MEP responses that are abolished by subsequent re-transection of the spinal cord in rats.

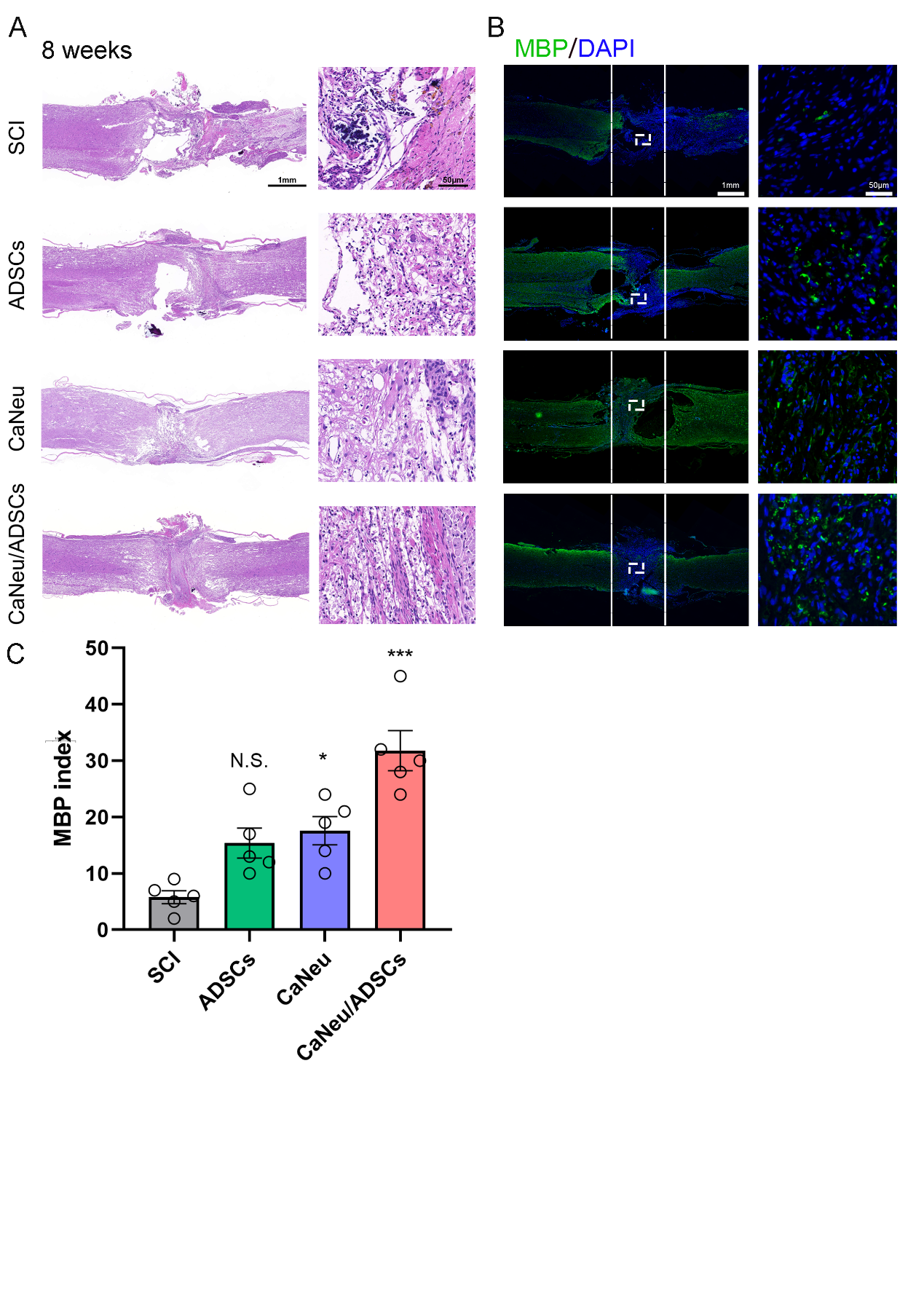

**Figure S8. CaNeu laden ADSCs promote remyelination after acute traumatic SCI.** (A) Representative H&E images of the injured spinal cord eight weeks after the administration of ADSCs, CaNeu and CaNeu/ADSCs, higher magnification views in right column. (B) Immunofluorescence image of MBP in lesion center in SCI, ADSCs, CaNeu and CaNeu/ADSCs-treated rats at 8weeks after SCI. (C) Quantification of MBP intensity. Scale bars indicate 1 mm, 50 μm, respectively. N. S.: no significance, *, *** indicate *P* < 0.05, *P* < 0.001, respectively, by one-way ANOVA followed by Tukey’s post hoc analysis. Data are expressed as mean ± S.E.M. n = 5 per group.

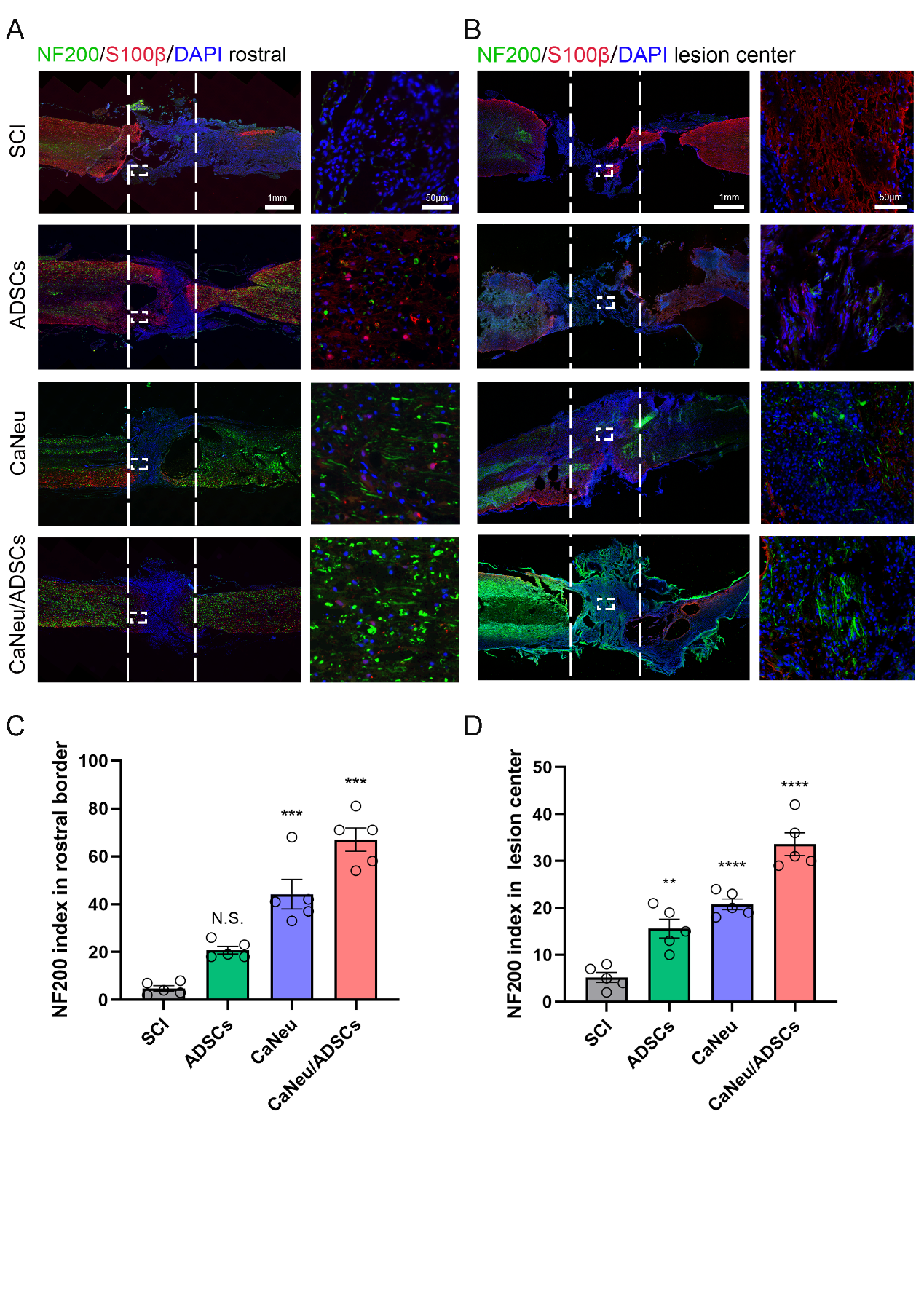

**Figure S9. CaNeu laden ADSCs promote axonal growth after acute traumatic SCI.** (A, B) Immunofluorescence image of NF200 in rostral border (A) and lesion center (B) in SCI, ADSCs, CaNeu and CaNeu/ADSCs-treated rats at 8 weeks after SCI. Scale bars indicate 1 mm, 50 μm, respectively. (C, D) Quantification of NF200 intensity in rostral border (C) and lesion center (D), respectively. N. S.: no significance, *, *** and **** indicate P < 0.05, P < 0.001 and P < 0.0001, respectively, by one-way ANOVA followed by Tukey’s post hoc analysis. Data are expressed as mean ± S.E.M. n = 5 per group.

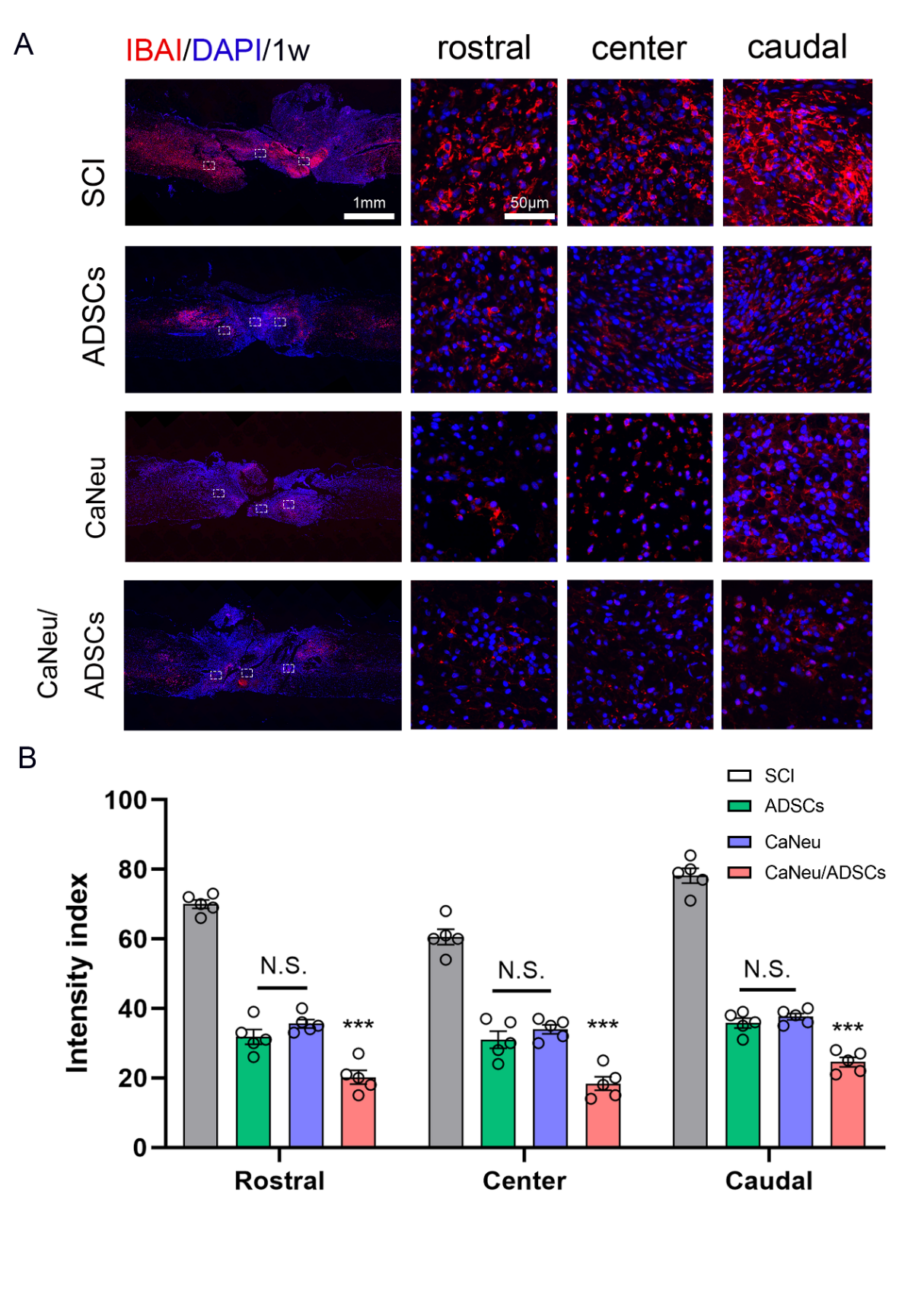

**Figure S10.** **Immune response** **of spinal cord sections at the implant site.** (A) Immunoﬂuorescence images showed that the percentage of IBA1^+^ cells in rostral/center/caudal of CaNeu/ADSCs group, CaNeu, ADSCs, and SCI groups per visual ﬁeld at 1 week after SCI. Scale bars indicate 1 mm, 50 μm, respectively. (B) Quantiﬁcation of IBA1 intensity. Error bars represent mean ± S.E.M. N. S. indicates no significance, *** indicates *P* < 0.001. Tukey's multiple comparisons test of two-way ANOVA. n = 5 replicate experiments per group.

**Table S1. Animals sacrificed in this study**

| Time points | Animal numbers/per group | Experiments |
| --- | --- | --- |
| 1w | 3 (all groups except sham) | IF staining |
| 1w | 3 (all groups) | WB |
| 2w | 3 (all groups except sham) | IF staining |
| 8w | 5 (all groups) | EM |
| 8w | 5 (all groups) | Electrophysiology/IF staining |
